# Squamous trans-differentiation of pancreatic cancer cells promotes stromal inflammation

**DOI:** 10.1101/833046

**Authors:** Tim D.D. Somerville, Giulia Biffi, Juliane Daßler-Plenker, Koji Miyabayashi, Yali Xu, Diogo Maia-Silva, Olaf Klingbeil, Osama E. Demerdash, Mikala Egeblad, David A. Tuveson, Christopher R. Vakoc

## Abstract

A highly aggressive subset of pancreatic ductal adenocarcinomas undergo trans-differentiation into the squamous lineage during disease progression. While the tumorigenic consequences of this aberrant cell fate transition are poorly understood, recent studies have identified a role for the master regulator TP63 in this process. Here, we investigated whether squamous trans-differentiation of pancreatic cancer cells can influence the phenotype of non-neoplastic cells in the tumor microenvironment. Conditioned media experiments revealed that squamous-subtype pancreatic cancer cells secrete factors that convert quiescent pancreatic stellate cells into a specialized subtype of cancer-associated fibroblasts (CAFs) that express inflammatory genes at high levels. We use gain- and loss-of-function approaches *in vivo* to show that squamous-subtype pancreatic tumor models become enriched with inflammatory CAFs and neutrophils in a TP63-dependent manner. These non cell-autonomous effects occur, at least in part, through TP63-mediated activation of enhancers at pro-inflammatory cytokine loci, which includes *IL1A* as a key target. Taken together, our findings reveal enhanced tissue inflammation as a consequence of squamous trans-differentiation in pancreatic cancer, thus highlighting an instructive role of tumor cell lineage in reprogramming the stromal microenvironment.

## INTRODUCTION

Pancreatic ductal adenocarcinoma (PDA) is one of the most lethal human tumors, with a five year survival rate below 10% (Siegel et al., 2019). Despite the overall association with poor clinical outcomes, a striking heterogeneity in the presentation and progression of this disease exists between individuals, including differing rates of metastatic spread and responses to cytotoxic chemotherapy (Collisson et al., 2019). Owing to this issue, a major objective in PDA research is to uncover molecular mechanisms that underpin this clinical heterogeneity. Such efforts may yield much-needed biomarkers capable of predicting disease progression, as well as personalized therapeutic strategies that target critical disease drivers in each patient’s tumor.

It has long been recognized that a subset of pancreatic tumors exhibit an ‘adenosquamous’ histology, which is characterized by the presence of both glandular and squamous neoplastic cells within the same tumor (Ishikawa et al., 1980; Morohoshi et al., 1983; Motojima et al., 1992). The designation of adenosquamous pancreatic cancer as a disease entity has been reinforced by several recent transcriptome profiling studies of human pancreatic tumors, which identified aberrant expression of squamous (also known as basal) lineage markers in ∼15% of samples, in association with exceptionally poor clinical outcomes (Bailey et al., 2016; Cancer Genome Atlas Research Network, 2017; Moffitt et al., 2015). While squamous cells are a normal cell type in several stratified epithelial tissues (e.g. skin, esophagus), they are not known to be present in the normal human pancreas (Basturk et al., 2005). This suggests an aberrant ductal-to-squamous epithelial cell fate transition induced during pancreatic tumorigenesis.

The functional relevance of squamous trans-differentiation in pancreatic cancer was unclear until recently, when studies from several laboratories, including our own, demonstrated that the transcription factor TP63 (delta N isoform, hereafter referred to as TP63 for simplicity) is the master regulator of the adenosquamous phenotype in PDA (Andricovich et al., 2018; Hamdan and Johnsen, 2018; Somerville et al., 2018). Once expressed, TP63 binds to thousands of genomic sites to nucleate the formation of active enhancers to drive expression of squamous lineage genes (e.g. *KRT5/6* and *S100A2*) in PDA cells (Hamdan and Johnsen, 2018; Somerville et al., 2018). Importantly, TP63 is both necessary and sufficient to endow PDA cells with the same squamous lineage transcriptional program that is observed in pancreatic tumors with adenosquamous histology (Bailey et al., 2016; Hamdan and Johnsen, 2018; Somerville et al., 2018). While the mechanisms that induce TP63 expression in PDA remain unclear, it has been shown in mice that inactivation of the tumor suppressor *Kdm6a* or over-expression of *Myc* can predispose pancreatic tumors to express squamous markers, albeit with partial penetrance (Andricovich et al., 2018; Witkiewicz et al., 2015). In support of the functional impact of squamous trans-differentiation in PDA, induction of TP63 leads to a collection of phenotypic alterations, including enhanced motility, invasion, and resistance to cytotoxic chemotherapy (Danilov et al., 2011; Somerville et al., 2018).

Evidence from human patients and mouse models supports a powerful effect of inflammation in driving PDA progression (Guerra et al., 2011; Guerra et al., 2007; Yadav and Lowenfels, 2013). A key step in this process is an elaboration of pro-inflammatory cytokines by tumor cells and non-neoplastic cells in the stromal compartment (Mantovani et al., 2008). For example, many pancreatic tumors are infiltrated with neutrophils, which suppress anti-tumor T-cell immunity in PDA mouse models and correlate with aggressive disease in humans (Bayne et al., 2012; Chao et al., 2016; Inoue et al., 2014; Shen et al., 2014; Steele et al., 2016). A major cell type in the stroma of PDA tumors are CAFs, which have historically been considered tumor-promoting, however recent studies suggest they may also have tumor restraining functions (Öhlund et al., 2014; Özdemir et al., 2014; Rhim et al., 2014). A subset of CAFs with myofibroblastic properties, termed myCAFs, appear to be involved in the production of extracellular matrix which limits drug delivery to the tumor (Olive et al., 2009; Provenzano et al., 2012; Sherman et al., 2014). Another subset of CAFs, termed iCAFs, have low expression of myCAF markers and instead produce high levels of inflammatory cytokines, such as IL-6, CXCL1, and LIF (Öhlund et al., 2017). Although the iCAF/myCAF ratio, the extent of neutrophil infiltration, and the overall level of stromal inflammation can vary significantly between PDA patients, the underlying features of cancer cells that drive these differences are only beginning to be understood (Vennin et al., 2019).

Here we provide evidence that TP63-expressing squamous PDA cells have an enhanced capability to promote inflammatory changes within the tumor microenvironment when compared to TP63-negative PDA cells. These findings are supported by *in vitro* and *in vivo* experiments, and include activation of an inflammatory transcriptional program in CAFs of the tumor stroma. These effects are mediated via a specific cytokine secretion phenotype of PDA cells downstream of TP63-mediated enhancer reprogramming. Taken together, these findings highlight how lineage alterations within cancer cells orchestrate non cell-autonomous changes in the tumor microenvironment.

## RESULTS

### A secretory phenotype of TP63-positive PDA cells that promotes inflammatory gene expression changes in CAFs *in vitro*

In a prior study, we found that ectopic expression of TP63 in the human PDA cell line SUIT2 (which lacks endogenous TP63 expression) enhanced cell growth when implanted into the mouse pancreas *in vivo*, yet this phenotype was absent under tissue culture conditions (Somerville et al., 2018). These findings led us to hypothesize that TP63 expression alters how PDA cells communicate with non-neoplastic cells in the tumor stroma. Here, we focused our studies on CAFs, given their established roles in PDA pathogenesis (Kalluri, 2016). To this end, we collected conditioned media from 12 human PDA cell lines exhibiting different expression levels of squamous markers (e.g. *TP63* and *KRT5*) (Fig. 1A-C and S1A). Of note, MIAPaca2 cells only express the TA isoform of TP63 (Fig. S1A) and lack expression of squamous lineage markers (Somerville et al., 2018) and in fact are more representative of a cell type of neuroendocrine origin (Yu et al., 2019). The collected media was then applied to quiescent murine pancreatic stellate cells (PSCs), which are a precursor of CAFs in PDA, and RNA-sequencing analysis was performed on the PSC cultures following 96 hours to evaluate the cellular response (Fig. 1D). An unsupervised clustering of the global transcriptional profile of the PSCs revealed three distinct groups (Fig. 1E). Two of the treated PSC cultures clustered closely with the untreated controls, suggesting they remained in a quiescent transcriptional state (Fig. 1E). However, two major groups were found to cluster away from the control PSC cultures and were termed Group 1 and Group 2 (Fig. 1E). Remarkably, the PSCs within the Group 2 cluster were all treated with conditioned media derived from the three PDA cell lines expressing TP63 (BxPC3, T3M4, and KLM1; Fig. 1A-B and S1A). In addition to this transcriptional phenotype, the conditioned media from TP63-expressing lines also led to a stronger induction of PSC proliferation than conditioned media collected from TP63-negative lines (Fig. S1B-C). We next extracted the subset of genes that discriminate the Group 1 and Group 2 clusters of PSC transcriptional responses, identifying 144 and 259 differentially expressed genes, respectively (Fig. 1F and Table S1). A Gene Set Enrichment Analysis (GSEA) of the Group 2 cluster revealed several top-ranking gene signatures associated with inflammation (Fig. 1G and S1D). Using transcriptome data from our prior study in which we identified iCAFs and myCAFs (Öhlund et al., 2017), we defined gene signatures associated with these two cell fates (Table S2) and observed a significant enrichment of the iCAF signature within the Group 2 set of PSCs (Fig. 1H). In contrast, the myCAF signature was significantly enriched within Group 1 PSCs (Fig. 1I). RT-qPCR analysis of human PSCs treated with conditioned media from human PDA cell lines further validated the correlation between TP63 expression in the cancer cells with iCAF induction (Fig. S1E). These data suggest that PDA cells harboring the squamous transcriptional profile secrete factors that convert PSCs into iCAFs *in vitro*.

**Figure 1.**
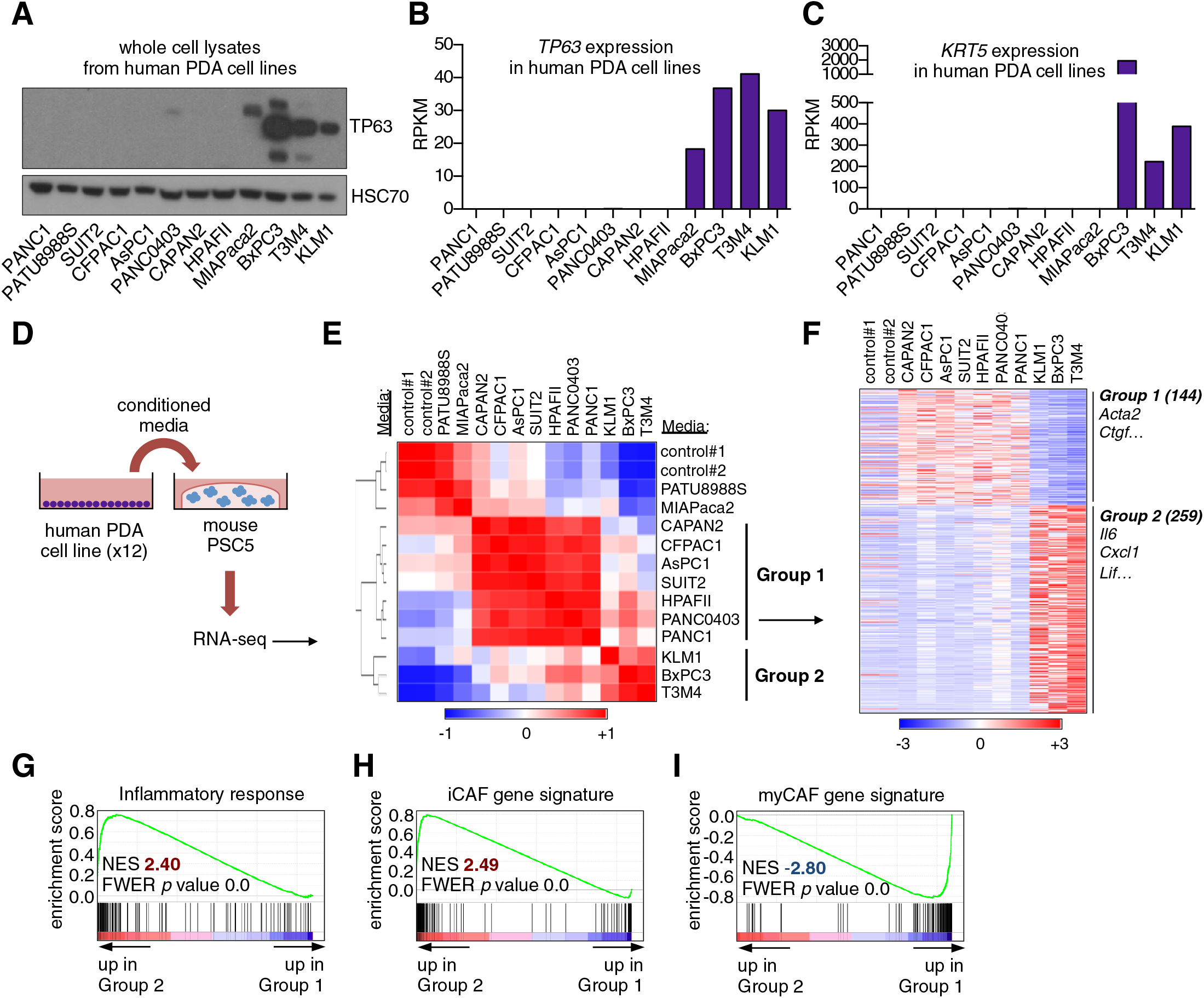
A secretory phenotype of TP63-positive PDA cells that promotes inflammatory gene expression changes in CAFs *in vitro*. (A) Western blot analysis showing TP63 expression in a panel of 12 human PDA cell lines. (B-C) Bar charts show expression of *TP63* and *KRT5* cell lines shown in (A). (D) Schematic of experimental workflow for RNA-seq analysis of PSCs following culture in Matrigel with conditioned from the 12 human PDA cell lines. (E) Heatmap representation of unsupervised hierarchical clustering of mouse PSCs based on their global transcriptional profile. Scale bar indicates Pearson correlation coefficient. Control refers to control media which was DMEM supplemented with 5% FBS. (F) Heatmap representation of differentially expressed genes from PSCs in Group 1 and Group 2. Selected genes in each group are listed. Scale bar indicates standardized expression value. (G) GSEA plot evaluating Hallmark Inflammatory Response genes based on their expression in Group 2 versus Group 1 cultures. (H-I) GSEA plots evaluating the iCAF and myCAF gene signatures in Group 2 versus Group 1 cultures. See also Figure S1.

### TP63 expression in PDA cells drives a secretory phenotype that induces iCAF formation *in vitro*

Based on the correlations described above, we next set out to determine the causality between TP63 and the iCAF-inducing secretory phenotype. To this end, we cultured two independent mouse PSC lines with conditioned media harvested from SUIT2-empty and SUIT2-TP63 cells and performed RT-qPCR analysis of iCAF marker genes in the PSC cultures (Fig. 2A). We observed a marked increase in iCAF markers (*Il6*, *Cxcl1*, and *Lif*) and a concomitant reduction of myCAF markers (*Acta2* and *Ctgf*) when the PSCs were cultured in the SUIT2-TP63 media compared to those cultured in SUIT2-empty media (Fig. 2B-C, S2A-B). In accord with the findings described above, mouse PSCs cultured in the SUIT2-TP63 conditioned media were more proliferative when compared to their counterparts cultured in SUIT2-empty conditioned media (Fig. S2C-D). Similar RT-qPCR experiments using human PSC cultures or using a metastatic mM1 organoid line, which was isolated from the KPC (*Kras^+/LSL-G12D^; Trp53^+/LSL-R172H^; Pdx1-Cre*) mouse model (Boj et al., 2015), ectopically expressing TP63 support the generality of these findings (Fig. 2D-E, S2E-F).

**Figure 2.**
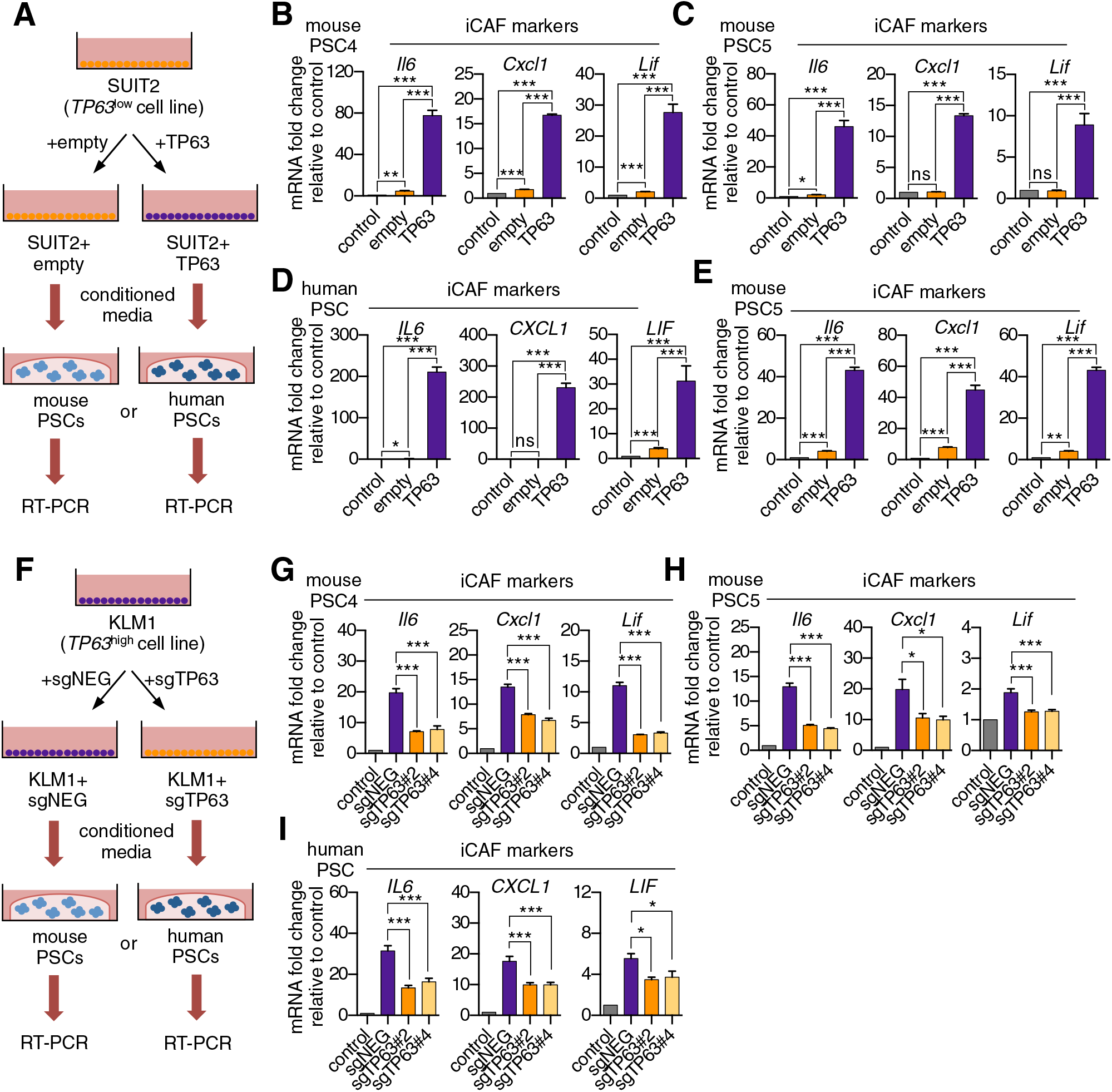
TP63 expression in PDA cells drives a secretory phenotype that induces iCAF formation *in vitro*. (A) Schematic of workflow for generating conditioned media from SUIT2-empty and SUIT2-TP63 cells and subsequent analysis of PSC cultures. (B-C) Bar charts showing RT-qPCR analysis for iCAF markers (*Il6*, *Cxcl1*, *Lif*) following culture of 2 mouse PSC lines in Matrigel for four days with the indicated conditioned media. (D) RT-qPCR analysis for the iCAF markers (*IL6*, *CXCL1*, *LIF*) following culture of human PSCs in Matrigel for four days with the indicated conditioned media. (E) Mouse mM1 organoids were infected with a TP63 cDNA or empty vector control. Bar chart shows RT-qPCR analysis for iCAF markers (*Il6*, *Cxcl1*, *Lif*) in the indicated conditions. Mean+SEM is shown. n=3. (F) Schematic of workflow for generating conditioned media from KLM1-Cas9 cells infected with TP63 sgRNAs (sgTP63#2 and sgTP63#4) or a control (sgNEG) and subsequent analysis of PSC cultures. (G-H) Bar chart showing RT-qPCR analysis for iCAF markers (*Il6*, *Cxcl1*, *Lif*) following culture of mouse and human PSCs in Matrigel with the indicated conditioned media. (I) RT-qPCR analysis for iCAF markers (*IL6*, *CXCL1*, *LIF*) following culture of human PSCs in Matrigel for four days with the indicated conditioned media. For all experiments control media represents DMEM supplemented with 5% FBS. Mean+SEM is shown. n=3. For B-E, \*\*\**p* <0.001, \*\**p* <0.01, \**p* <0.05 by Student’s t-test and for G-I, \*\*\**p* <0.001, \*\**p* <0.01, \**p* <0.05 by one-way ANOVA with Dunnett’s test for multiple comparisons. See also Figure S2.

We next performed loss-of-function studies using conditioned media from KLM1-Cas9 cells infected with two independent sgRNAs targeting TP63 or a control sgRNA (Fig. 2F). Notably, KLM1 is the only TP63-postive PDA cell line we have identified which does not exhibit a growth arrest phenotype *in vitro* following TP63 inactivation (Fig. S2G-H) (Somerville et al., 2018). Importantly, KLM1 cells harbor a similar transcriptional and epigenomic profile as observed in other TP63-expressing PDA cell lines (Fig. S2I-K). In addition, KLM1 cells also induce a similar iCAF phenotype as seen with other TP63-expressing lines (Fig. 1E-F). Hence, this cell line is useful for performing TP63 loss-of-function experiments evaluating for secretory phenotypes without the confounding effect of impairing cancer cell fitness.

While conditioned media from KLM1 cells led to a significant induction of iCAF markers and repression of myCAF markers in mouse and human PSCs, this effect was significantly attenuated using TP63-knockout KLM1 cells (Fig. 2G-I and S2L-N). In addition, the proliferation phenotype of the conditioned media-treated PSCs was also attenuated following TP63 inactivation (Fig. S2O-P). Taken together, these experiments demonstrate the critical role of TP63 in regulating the secretory phenotype in squamous-like models of PDA, which leads to the induction of inflammation-associated transcriptional changes in CAFs.

### Ectopic expression of TP63 in PDA cells promotes inflammation-associated transcriptional changes in the tumor microenvironment

We next evaluated the relevance of the TP63-induced secretory phenotype in an *in vivo* model of PDA. For this purpose, we first infected SUIT2-luciferase cells (Somerville et al., 2018) with a TP63 cDNA or empty vector prior to orthotopic transplantation into the pancreas of NOD-scid gamma (NSG) mice. In accordance with our previous findings (Somerville et al., 2018), TP63 expression enhanced the growth of SUIT2 cells *in vivo*, but not *in vitro* (Fig. 3A-B, S3A-D). Importantly, we validated the *in vivo*-specific growth advantage caused by ectopic TP63 expression in the mM1 organoids following transplantation into the pancreas of C57BL/6 mice (Fig. S3E-K). Tumors from the SUIT2 xenograft model were harvested and subjected to bulk RNA-seq analysis. Mapping of human and mouse transcripts to their respective genomes allowed us to discriminate the transcriptional changes happening in the human cancer cells from those of non-neoplastic mouse cells comprising the tumor stroma (Fig. 3C).

**Figure 3.**
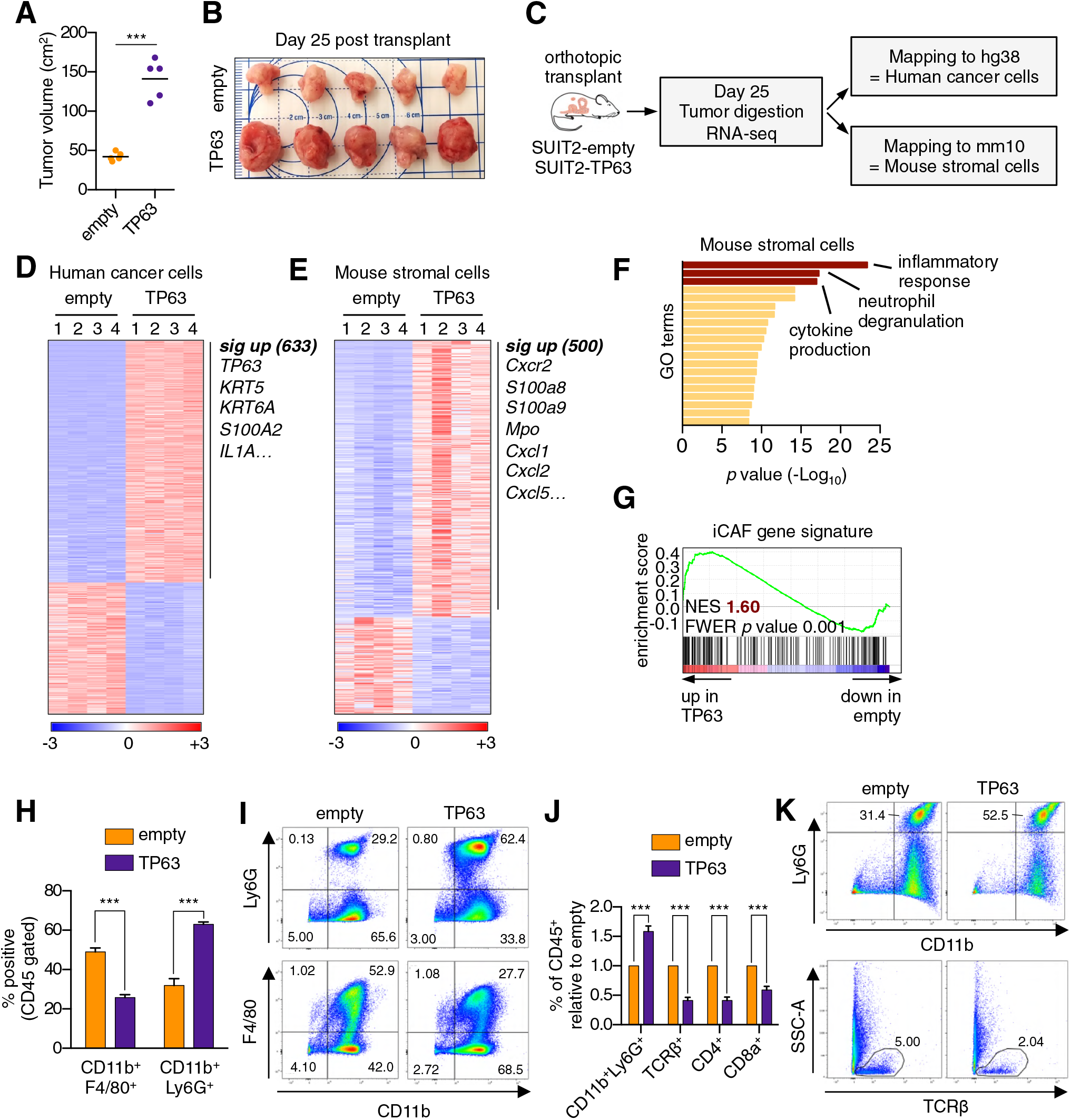
Ectopic expression of TP63 in PDA cells promotes inflammation-associated transcriptional changes in the tumor microenvironment. (A-I) SUIT2-empty and SUIT2-TP63 PDA cells harboring a luciferase transgene were transplanted to the pancreas of NSG mice. (A-B) Quantification of tumor volume (A) and images of tumors (B) on day 25 post-transplantation. (C) Experimental workflow for RNA-seq analysis of bulk tumor tissue. (D-E) Heatmap representation of differentially expressed genes that unambiguously map to (D) the human genome and therefore are derived from human cancer cells or (E) the mouse genome and therefore are derived from mouse stromal cells. Selected genes in each group are listed. Scale bar indicates standardized expression value. (F) Ontology analysis of the 500 significantly up-regulated mouse genes using Metascape. GO terms are ranked by their significance (*p* value) and the most significant terms (-log_10_ *p* value >15) are highlighted. (G) GSEA plot evaluating the iCAF signature in the mouse stromal compartment of SUIT2-TP63 versus SUIT2-empty tumors. (H) Quantification of flow cytometry analysis on bulk tumor tissues from SUIT2 xenografts. n=5 mice per group. \*\*\**p* <0.001 by Student’s t-test. (I) Representative flow cytometry plots from (H). (J-K) KPC mM1 organoids with a TP63 cDNA or empty vector control and transplanted to the pancreas of C57BL6 mice. (J) Quantification of flow cytometry analysis of CD45^+^ cells in the tumors. n=4-5 mice per group. \*\*\**p* <0.001 by Student’s t-test. (K) Representative flow cytometry plots from (J). See also Figure S3.

In the human cancer cell compartment, we identified 633 genes that were significantly upregulated in SUIT2-TP63 tumors compared to controls, which largely corresponded to squamous lineage TP63 target genes (e.g. *S100A2*, *KRT5)* (Fig. 3D, S3L and Table S3) (Bailey et al., 2016; Somerville et al., 2018). Within the mouse stromal compartment, we identified 500 genes that were significantly upregulated in SUIT2-TP63 tumors compared to controls (Fig. 3E and Table S3). An ontology analysis of this set of genes revealed enrichment for inflammatory responses, neutrophil degranulation, and cytokine production as top-ranking gene sets (Fig. 3F). Importantly, the transcriptional signature of iCAFs was significantly more enriched in the mouse stromal compartment of SUIT2-TP63 relative to SUIT2-empty tumors, which is consistent with our conditioned media experiments (Fig. 3G). Fluorescent activated cell sorting (FACS) of fibroblasts from these two groups of tumors confirmed an increase in iCAF markers and a decrease in myCAF markers in the SUIT2-TP63 tumors (Fig. S3M-O). Flow cytometry analysis confirmed that SUIT2-TP63 tumors were more infiltrated with neutrophils when compared to control tumors, which is a marker of enhanced tissue inflammation (Fig. 3H-I). A similar neutrophil infiltration phenotype was observed in the orthotopic mM1-TP63 tumors, which also correlated with a decrease in infiltrating CD4 and CD8 T cells (Fig. 3J-K). This finding is consistent with prior evidence that tumor-associated neutrophils have an immunosuppressive effect in PDA (Bayne et al., 2012; Chao et al., 2016). Finally, plasma concentrations of the cytokine IL-6, a marker of systemic inflammation (Nishimoto and Kishimoto, 2006), were ∼10-fold higher in the mice bearing SUIT2-TP63 versus SUIT2-empty tumors (Fig. S3P). Collectively, these experiments suggest that TP63-positive PDA cells can induce inflammatory changes in the tumor microenvironment *in vivo*.

### Knockout of TP63 in an orthotopic PDA tumor model attenuates the inflammatory signature of fibroblasts and immune cells

We next addressed the impact of TP63 inactivation on the tumor microenvironment *in vivo*. To this end, KLM1-Cas9-luciferase cells were transduced with TP63 sgRNAs before transplantation into the pancreas of NSG mice. By monitoring tumor progression using bioluminescent imaging, we found that TP63 inactivation resulted in a significant reduction in tumor growth when compared to control cells (Fig. 4A-B), which is consistent with *in vivo*-specific growth enhancement caused by acute TP63 expression in SUIT2 and mM1 orthotopic models. In order to study the impact of TP63 inactivation on the tumor microenvironment while controlling for overall tumor burden, we allowed sufficient time for tumors to form in each group before harvesting for analysis when a critical bioluminescent signal was reached. By this endpoint metric, cells transduced with the two independent TP63 sgRNAs formed tumors with a significantly longer latency versus controls (Fig. 4C). Immunohistochemical analysis for TP63 revealed an ‘escaper’ group (sgTP63#2) that gave rise to TP63-positive tumors, suggesting the emergence of TP63-positive clones within the pooled population of transplanted cells (Fig. 4D and S4A). However, in the second cohort where TP63 was inactivated (sgTP63#4), we identified largely TP63-negative tumors at endpoint, allowing us to interrogate the tumor microenvironment in a loss of function context (Fig. 4D-E and S4B).

**Figure 4.**
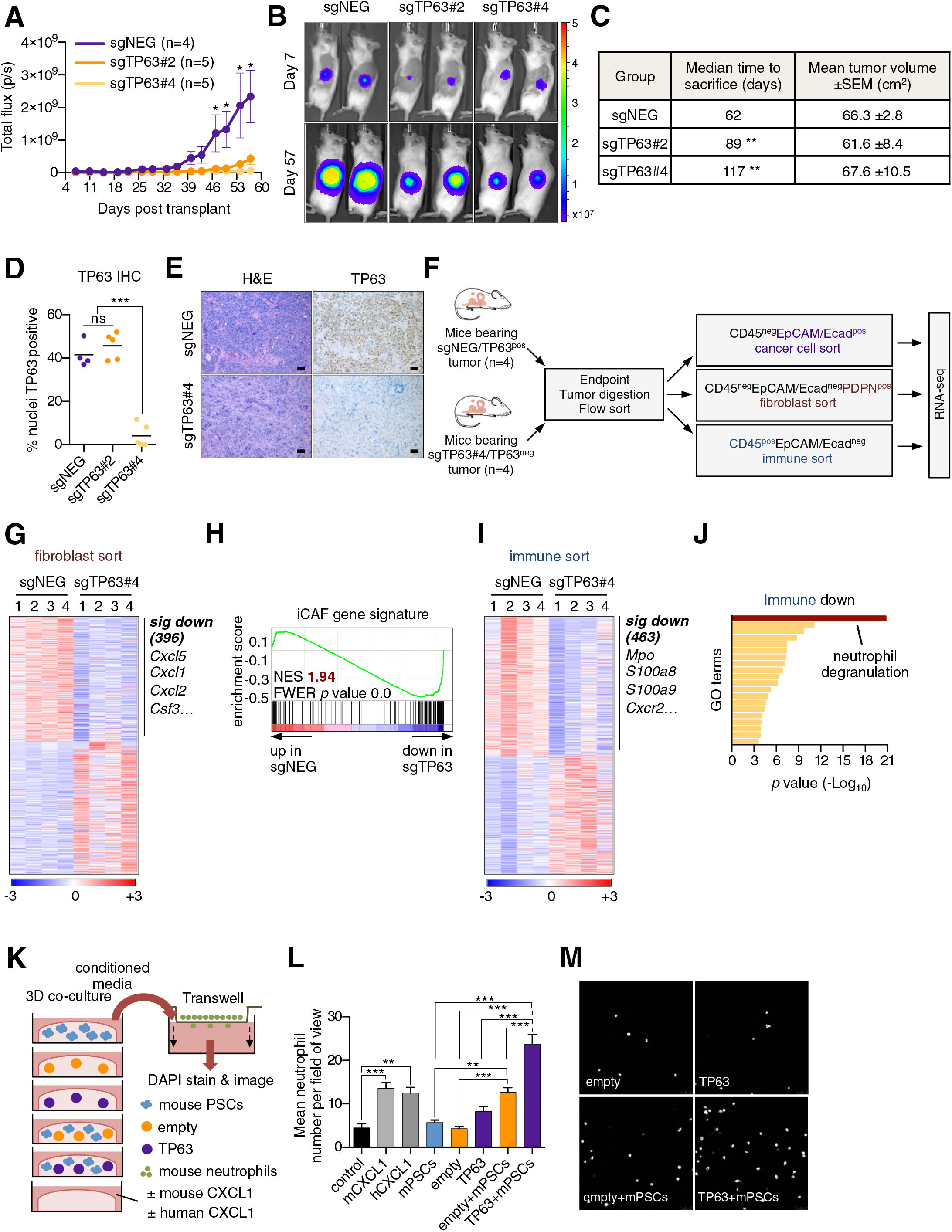
Knockout of TP63 in an orthotopic PDA tumor model attenuates the inflammatory signature of fibroblasts and immune cells. (A-E) KLM1-Cas9 cells expressing a luciferase transgene were infected with two independent TP63 or control (sgNEG) sgRNAs and transplanted to the pancreas of NSG mice. (A) Quantification of bioluminescence signal. Mean±SEM is shown. Mice were imaged every four days between day seven and 57 post-transplantation. n = 4-5 mice per group. \**p* <0.05 by two-way ANOVA with Sidak’s test for multiple comparisons. (B) Representative images at days 7 and 57 post transplant from (A). (C) Table summarizing the median time mice from each group were sacrificed and the tumor volume at endpoint. Endpoint was determined as a bioluminescence signal >3×10^9^ p/s for each individual mouse. \*\**p <*0.01 by log rank (Mantel-Cox) test. (D) Quantification of TP63 expression as determined by immunohistochemical staining of tumors from the indicated experimental groups. n=4-5 mice per group. \*\*\**p* <0.001 by one-way ANOVA with Tukey’s test for multiple comparisons. (E) Representative images from (D). Scale bar indicates 50µm. (F) Schematic of experimental workflow and sorting strategy for enriching for human cancer cells, mouse fibroblasts and mouse immune cells and subsequent RNA-seq analysis. (G-H) Heatmap representation of differentially expressed genes from the mouse fibroblast compartment (G) and immune compartment (H) and the indicated tumor samples. Selected genes in each group are listed. Scale bar indicates standardized expression value. (I) GSEA plot evaluating the iCAF signature in the mouse fibroblast compartment of TP63-positive versus control (sgNEG) tumors. (J) Ontology analysis of the 463 significantly down-regulated mouse genes from immune compartment using Metascape. GO terms are ranked by their significance (*p* value) and the most significant term (-log_10_ *p* value >15) is highlighted. (K-M) SUIT2-empty or SUIT2-TP63 cells were cultured in isolation or together with mouse PSCs in Matigrel for four days before conditioned media was harvested and used for a trans-well migration assay with freshly isolated mouse neutrophils. (K) Schematic of experimental workflow. (L) Quantification of trans-well neutrophil migration in the indicated media conditions as determined by confocal imaging. Mean+SEM is shown. n=3. \*\*\**p* <0.001, \*\**p* <0.01 by two-way ANOVA with Sidak’s test for multiple comparisons. All significant interactions are shown. (M) Representative confocal images from (L). See also Figure S4.

To achieve this, we FACS-purified human cancer cells, immune cells and fibroblasts and performed RNA-seq analysis (Fig. 4F). Within the cancer cell compartment, we identified 459 genes that were significantly downregulated in the TP63-knockout tumors (Table S4). As expected, this included *TP63* as well as its downstream target genes, *KRT5*, *KRT6A* and *S100A2* (Fig. S4C-D). Within the sorted fibroblast compartment, we identified 396 genes that were significantly downregulated in the TP63-knockout tumors, which included a significant suppression of the iCAF gene signature (Fig. 4G-H and Table S4). Finally, within the sorted immune cell fraction we identified 463 genes that were significantly down-regulated in the TP63-knockout tumors, which was enriched for markers of neutrophils, such as *Mpo*, *S100a8*, *S100a9* and *Cxcr2* (Fig. 4I-J and Table S4). Taken together, these findings demonstrate that inactivation of TP63 dampens the inflammation-associated transcriptional response present in the CAF and immune compartments of the stroma.

The findings above led us to investigate whether TP63-expressing PDA cells can collaborate with CAFs to increase neutrophil infiltration. In order to model this *in vitro*, we established a co-culture system in which human SUIT2-empty or SUIT2-TP63 cells were cultured in Matrigel either alone or in combination with mouse PSCs (Fig. 4K). RT-qPCR analysis for murine-specific transcripts confirmed that SUIT2-TP63 cells augmented the iCAF phenotype versus controls in this co-culture system (Fig. S4E). Of note, this included a marked upregulation of the iCAF gene *Cxcl1*, which encodes a powerful neutrophil chemoattractant (Moser et al., 1990). We found that the SUIT2-TP63/PSC co-cultures released soluble factors that increased the chemotaxis of primary mouse neutrophils and, importantly, this phenotype was enhanced in the SUIT2-TP63/PSC co-culture as compared to each cell type individually (Fig. 4L-M). Taken together, these experiments suggest that TP63-positive PDA cells can collaborate with CAFs to promote neutrophil migration.

### TP63 activates enhancer elements and transcription of genes encoding pro-inflammatory cytokines in PDA cells

We next sought to define a molecular mechanism by which TP63 drives a pro-inflammatory secretory phenotype by interrogating the transcriptional changes that occur following ectopic TP63 expression in SUIT2 cells (Somerville et al., 2018). GSEA comparing SUIT2-TP63 to SUIT2-empty cells revealed the activation of a number of inflammation-related pathways as top-ranking gene sets following TP63 expression (Fig. 5A-C and Fig. S5A). Of note, a number of these pathways overlap with those activated in iCAFs (Biffi et al., 2019). Next, we interrogated the transcriptional changes of inflammatory genes that occur following acute TP63 inactivation in BxPC3 cells, a human PDA cell line that expresses TP63 at high levels (Somerville et al., 2018). These combined analyses identified *IL1A*, *IL1B*, *CXCL1*, *CXCL8*, *CSF2* (*GM-CSF*), *CCL20*, *AREG*, *FJX1*, and *ADORA2B* as the top inflammation-related genes regulated by TP63 (Fig. 5C and Fig. S5B). Importantly, an analysis of TP63 over-expression and knockout/knockdown RNA-seq datasets performed in nine different human PDA lines or organoids (Somerville et al., 2018) revealed that this group of inflammatory genes is regulated by TP63 in these different contexts, albeit with a degree of heterogeneity (Fig. 5D and Fig. S5C). ChIP-seq analysis performed in BxPC3 cells revealed that each of these genes is located near a TP63-bound enhancer element, suggesting direct transcriptional regulation (Fig. S5D). Among these TP63 targets, *IL1A* and *IL1B* were outliers, as their expression was the most affected by TP63 perturbation (Fig. 5D). ChIP-seq analysis revealed a TP63-induced super-enhancer located at an intergenic region between the *IL1A* and *IL1B* genes (Fig. 5E and Fig. S5E-H). Using CRISPR-interference to target this region for inactivation (Qi et al., 2013), we verified that this group of TP63-bound enhancers was responsible for activating *IL1A* and *IL1B* expression (Fig. 5E, 5F and Fig. S5I). Taken together, these findings suggest that TP63 promotes enhancer-mediated transcriptional activation of genes encoding secreted factors with established roles in promoting inflammation.

**Figure 5.**
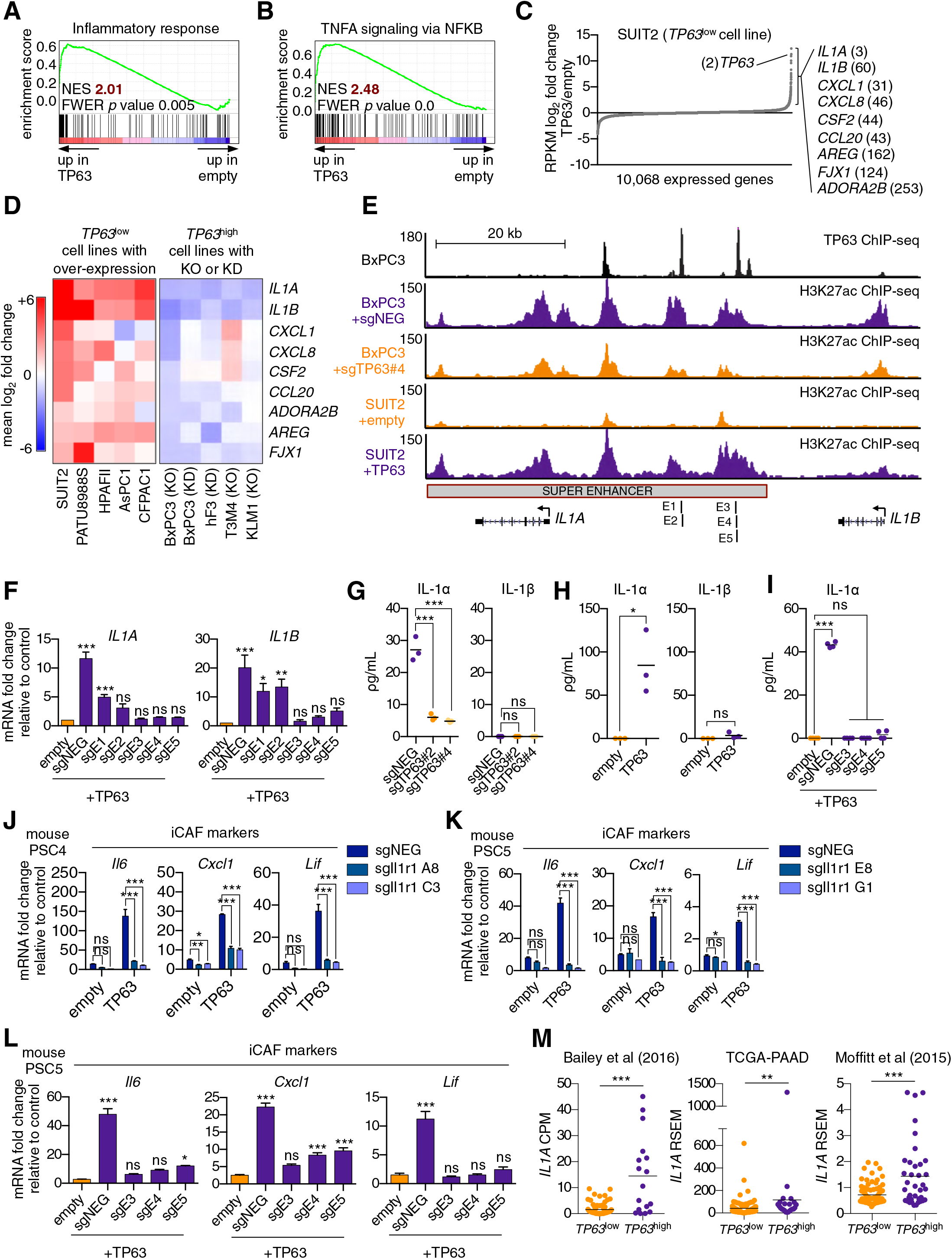
TP63 activates enhancer elements and transcription of genes encoding pro-inflammatory cytokines in PDA cells. (A-B) GSEA plots evaluating the indicated Hallmark gene signatures upon TP63 expression in SUIT2 cells. (C) Scatter plot shows the mean log_2_ fold change of expressed genes upon TP63 expression in SUIT2 cells. Genes with a mean log_2_ fold change >1 in this dataset and the BxPC3+sgTP63 dataset (Fig. S5B) that are also found in the gene signatures shown in A and B are highlighted along with the rank. Data are from Somerville et al (2018). (D) Heatmap shows gene expression changes in those genes shown in (C) in the indicated cell lines following TP63 over-expression (left panel) or knockout/knockdown (right panel). Scale bar indicates mean log_2_ fold change versus control. (E) ChIP-seq profiles of TP63 in BxPC3 cells (top track) and H3K27ac (bottom four tracks) following TP63 knockout in BxPC3 cells or overexpression in SUIT2 cells surrounding the *IL1* locus. The H3K27ac regions identified as a super enhancer by ROSE analysis is indicated along with the positions of sgRNAs targeting this enhancer used for experiments shown in (F) and (I). (F) RT-qPCR analysis for *IL1A* and *IL1B* in SUIT2-empty or SUIT2-TP63 cells infected with dCas9 fused with the KRAB repression domain and the indicated sgRNAs. The positions of the sgRNAs targeting the IL1 super enhancer are shown in (E). Mean+SEM is shown. n=3. \*\*\**p* <0.001, \*\**p* <0.01, \**p* <0.05 versus empty control by one-way ANOVA with Dunnett’s test for multiple comparisons. (G-H) ELISA for IL-1α and IL-1β from conditioned media harvested from KLM1-Cas9 cells infected with the indicated sgRNA (G) or SUIT2-empty and SUIT2-TP63 cells (H). (I) ELISA for IL-1α in SUIT2-empty or SUIT2-TP63 cells infected with dCas9 fused with the KRAB repression domain and the indicated sgRNAs. \*\*\**p* <0.001, \**p* <0.05 by one-way ANOVA with Tukey’s test for multiple comparisons. ns = not significant. (J-K) Bar charts showing RT-qPCR analysis for iCAF markers (*Il6*, *Cxcl1*, *Lif*) following culture of two mouse PSC lines from which the IL1 receptor was clonally knocked out with CRISPR in Matrigel for four days with conditioned media from SUIT2-empty or SUIT2-TP63 cells. (L) Bar charts showing RT-qPCR analysis for iCAF markers (*Il6*, *Cxcl1*, *Lif*) following culture of PSCs in Matrigel for four days with the conditioned media harvested from cells shown in (I). For all experiments control media represents DMEM supplemented with 5% FBS. Mean+SEM is shown. n=3. \*\*\**p* <0.001, \*\**p* <0.01, \**p* <0.05 by one-way ANOVA with Dunnett’s test for multiple comparisons. (M) Quantification of *IL1A* mRNA levels in the indicated studies. Each dot represents one patient sample and patients were stratified as *TP63*^high^ or *TP63*^low^ as described in Somerville et al (2018). ***p <0.001, **p <0.01 by Student’s t-test. ns, not significant. See also Figure S5.

### IL-1*α* is the key secreted molecule induced by TP63 in PDA cells that promotes iCAF induction

We focused our subsequent functional experiments on *IL1A* and *IL1B*, which respectively encode interleukin 1 alpha (IL-1α) and beta (IL-1β) and are important mediators of inflammatory responses (Di Paolo and Shayakhmetov, 2016; Dinarello and Wolff, 1993). We recently demonstrated a critical role for IL-1*α*in promoting the conversion of PSCs to iCAFs through the induction of JAK/STAT signaling (Biffi et al., 2019). In order to assess whether IL-1α and IL-1β are secreted by TP63-expressing PDA cells, we performed enzyme-linked immunosorbent assay (ELISA) experiments for these two proteins on conditioned media following TP63 knockout in KLM1 cells. This assay efficiently detected IL-1α, but not IL-1β, in the control setting and the levels of secreted IL-1α were significantly reduced by the two independent TP63 sgRNAs (Fig. 5G). An absence of IL-1β secretion in this context might be due to a lack of inflammasome activation, which is a known requirement for IL-β, but not IL-*α* secretion (Di Paolo and Shayakhmetov, 2016). Consistent with this observation, ectopic expression of TP63 in SUIT2 cells led to robust secretion of IL-1α but not IL-1β and this effect could be ablated by CRISPR-interference-mediated targeting of the IL1 super-enhancer (Fig. 5H-I). Taken together, these results indicate that both *IL1A* and *IL1B* are regulated by TP63 at the transcriptional level, but only IL-1α is secreted at detectable levels by TP63-expressing PDA cells *in vitro*.

In order to investigate whether TP63 driven iCAF induction is conferred via an IL-1-mediated signaling mechanism, we used mouse PSC lines in which the IL1 receptor (Il1r1) was knocked out using CRISPR (Biffi et al., 2019) and performed experiments with conditioned media harvested from SUIT2-TP63 or SUIT2-empty cells. Consistent with the results above, PSCs infected with the control sgRNA displayed a marked increase in iCAF gene expression following culture in SUIT2-TP63 conditioned media. However, this induction was absent in the IL1 receptor knock out lines (Fig. 5J-K, S5J-K). We confirmed the dependency on IL1 signaling for TP63 driven iCAF induction by performing conditioned media experiments using SUIT2-TP63 cells following inactivation of the IL1 super-enhancer via CRISPR interference (Fig. 5L and Fig. S5L). These data demonstrate that TP63 expressing PDA cells induce an iCAF phenotype via paracrine IL-1α signaling.

Finally, in order to ascertain whether TP63 expression is associated with increased *IL1A* expression in human pancreatic cancer patients, we interrogated three independent datasets that have profiled the transcriptome of bulk tumor tissue (Bailey et al., 2016; Cancer Genome Atlas Research Network, 2017; Moffitt et al., 2015). In all three datasets, the expression of *IL1A* was found to be significantly elevated in *TP63*^high^ versus *TP63*^low^ patient samples (Fig. 5M).

## Discussion

Prior studies have revealed PDA to be a heterogeneous disease at the molecular level with respect to both the tumor and stromal compartments (Bailey et al., 2016; Cancer Genome Atlas Research Network, 2017; Collisson et al., 2011; Moffitt et al., 2015). As both compartments are comprised of cell types that display phenotypic plasticity (Öhlund et al., 2017; Yuan et al., 2019), dissecting the molecular cross-talk between cells that define each patient’s tumor remains a formidable challenge. Although prior studies have made correlations between epithelial and stromal transcriptomes in PDA (Maurer et al., 2019; Moffitt et al., 2015; Puleo et al., 2018), it remains unclear whether the evolution of these two compartments is dependent on one another.

Here, we have defined a mechanism by which tumor cells gain an ability to reprogram their local microenvironment through the acquisition of TP63 expression, which is a master regulator of the squamous subtype of PDA (Hamdan and Johnsen, 2018; Somerville et al., 2018). Mechanistically, we have shown that TP63 activates a pro-inflammatory secretory phenotype that triggers iCAF induction through an IL-1α-dependent signaling mechanism. Importantly, we have verified the correlation between *TP63* and *IL1A* expression in three independent transcriptome studies of human PDA tumors. In addition, a recent transcriptome study of human tissue samples revealed a correlation between squamous subtype PDA tumors and inflammation-associated changes in the stroma, which validates that our observations in experimental models are in accord with clinical observations (de Santiago et al., 2019). While the aberrant expression of TP63 confers a myriad of tumor cell-intrinsic phenotypes in PDA, such as enhanced motility, invasion, and resistance to cytotoxic chemotherapy (Danilov et al., 2011; Somerville et al., 2018), our work points to a powerful non cell-autonomous effect of TP63 in PDA in promoting inflammation.

We previously demonstrated that IL1/JAK/STAT signaling is the key pathway responsible for iCAF formation in PDA (Biffi et al., 2019). While a basal level of IL1 is expressed in PDA tumor cells that lack TP63 expression (Tjomsland et al., 2011; Zhang et al., 2018), a key finding in our current study is in demonstrating that a remarkable heterogeneity exists in IL1 expression in the human disease, with the highest levels being found in TP63-expressing squamous subtype tumors. We account for this observation by demonstrating that a super-enhancer is installed by TP63 at the *IL1* locus, which enables a >30-fold increase in IL-1α secretion to enhance iCAF formation. One expected outcome of TP63-mediated *IL1A* hyperactivation would be a strengthened concentration gradient of IL-1α emanating from tumor cells to promote iCAF formation at distal locations within the stroma. While the abundance and spatial localization of iCAFs and myCAFs are likely regulated by several signaling pathways (e.g. TGFβ), our *in vivo* findings suggest that the acquisition of TP63 expression in the tumor leads to a potent increase in IL-1α secretion to drive iCAF enrichment.

Many of the genes regulated by TP63 in PDA cells correspond to those activated by the pro-inflammatory transcription factor NFκB. Indeed, it has been previously shown that TP63 and NFκB co-occupy a common set of target genes in head and neck squamous cell carcinoma, and this is associated with enhanced inflammation *in vivo* (Yang et al., 2011). In addition, NFκB is often constitutively active in most pancreatic cancer tissues and confers resistance to apoptosis (Dong et al., 2002; Wang et al., 1999). In this context, constitutive activation of NFκB requires an IL-1α-driven feed-forward amplification loop to promote PDA development (Ling et al., 2012; Niu et al., 2004). Maintenance of an inflamed CAF state has also been shown to be dependent on the NFκB signaling pathway (Biffi et al., 2019; Erez et al., 2010). Taken together, these studies raise the possibility that TP63 functions as an amplifier of NFkB transcriptional output to drive enhanced secretion of inflammatory mediators in squamous subtype PDA tumors. In addition to IL-1α, TP63 promotes expression of other pro-inflammatory cytokines that have been implicated in the development and progression of PDA, such as CXCL1 and GM-CSF (Bayne et al., 2012; Li et al., 2018; Pylayeva-Gupta et al., 2012). Notably, both of these cytokines are also produced by iCAFs, suggesting that the pro-tumorigenic effects of these factors would be amplified by IL-1α-mediated crosstalk between TP63-expressing tumor cells and iCAFs.

An association between squamous cell carcinomas and infiltrating inflammatory cells has been observed previously in other tumor contexts (Andreu et al., 2010; Coussens et al., 1999; Erez et al., 2010; Eruslanov et al., 2014; Ferone et al., 2016; Kargl et al., 2017; Xu et al., 2014; Yang et al., 2011). In one study, squamous trans-differentiation in a lung adenocarcinoma mouse model was associated with neutrophil infiltration and other markers of inflammation (Mollaoglu et al., 2018). In this context, inflammatory changes were promoted by the squamous lineage transcription factor SOX2 through activation of CXCL5 expression (Mollaoglu et al., 2018). Taken together with our findings, these results suggest that inflammation is more generally associated with squamous lineage tumors through multiple distinct mechanisms. Interestingly, a major function of the normal squamous epithelium is to serve as a protective barrier to exogenous insults. As such, inflammatory pathways must be tightly regulated and poised for a rapid response in these tissues. Notably, squamous cells of the epidermis are known to express large amounts of IL-1α at steady state (Hauser et al., 1986) and *IL1A* is know to be a key TP63 target in normal human keratinocytes (Barton et al., 2010). For this reason, amplified inflammation may be an inevitable consequence of tumor trans-differentiation into the squamous lineage irrespective of tissue context.

Histopathological definitions of PDA are often arbitrary and imprecise, and offer limited clinical utility with respect to patient management and treatment selection (Klöppel and Luettges, 2001). Adenosquamous carcinoma, for example, is defined by those tumors exhibiting >15% squamous differentiation and is considered an uncommon variant of PDA. Studies that have identified molecular subtypes of PDA using transcriptomic analyses provide the opportunity to define a new molecular taxonomy for pancreatic cancer that is more clinically applicable (Collisson et al., 2019). The challenge now is to understand the underlying biology that renders these molecular subtypes phenotypically and functionally distinct and to identify robust markers for prospective patient stratification. The findings presented in our study point to TP63 as a functional biomarker that identifies a poor prognostic subgroup of PDA patients in which inflammatory pathways are amplified. As such, these patients may benefit from targeted therapies that aim to dampen the inflammatory response. For example, monoclonal antibodies targeting IL-1α or IL-β have shown promise as potential cancer therapies (Hickish et al., 2017; Hong et al., 2014; Ridker et al., 2017). The findings presented in this study would suggest that such agents may show efficacy in appropriately stratified PDA patients.

## MATERIALS AND METHODS

### Cell Lines and Cell Culture

Mouse PSCs and pancreatic organoid lines were previously described (Boj et al., 2015; Öhlund et al., 2017). Human PSCs were purchased from ScienCell (3830). Mouse PSCs in which the Il1r1 was knocked out using CRISPR were previously described (Biffi et al., 2019). Mouse and human PSCs were cultured in DMEM (10-013-CV; Fisher Scientific) containing 5% FBS. HEK 293T cells were cultured in DMEM containing 10% FBS. Human PDA cell lines were cultured in RPMI 1640 (Gibco) containing 10% FBS. All cells were cultured with 1% L-glutamine and 1% Penicillin/Streptomycin at 37°C with 5% CO_2_. All cell lines were routinely tested for mycoplasma. For conditioned media experiments, human PDA cell lines or mouse organoids were cultured for 3-4 days in DMEM containing 5% FBS before media was collected and filtered through a 0.45µm syringe filter to remove debris. Conditioned media was stored at 4°C for no more than 3-4 days or snap frozen and stored at −80°C prior to use.

### Lentiviral Production and Infection

Lentivirus was produced in HEK 293T cells by transfecting plasmid DNA and packaging plasmids (VSVG and psPAX2) using Polyethylenimine (PEI 25000; Polysciences; Cat# 23966-1). Media was replaced with target media 6-8 hours following transfection and lentivirus-containing supernatant was subsequently collected every 12 hours for 48 hours prior to filtration through a 0.45µm filter. For infection of human PDA cells, cell suspensions were mixed with lentiviral-containing supernatant supplemented with polybreane to a final concentration of 4µg/ml. Cells were plated in tissue culture plates of the appropriate size and lentiviral-containing supernatant was replaced with fresh media after an incubation period of 24 hours. For infection of pancreatic organoids, lentivirus was first concentrated 10x in mouse organoid media (Boj et al., 2015) using Lenti-X^TM^ concentrator according to the manufacturer’s instructions. Organoid cultures were then dissociated into single cells, and spinoculated by centrifugation as described previously (Roe et al., 2017).

### CRISPR-based Targeting

To generate cell lines in which TP63 had been stably knocked out, KLM1 cells expressing Cas9 in the LentiV-Cas9-puro vector (addgene # 108100) were infected in a pooled fashion with control or TP63 sgRNAs in the LRNG vector (Roe et al., 2017). Two days post infection with sgRNAs, transduced cells were selected with 1mg/ml of G418 for five days before they were used for conditioned media experiments or orthotopic transplantations. For GFP-depletion assays, KLM1 and T3M4 cells were infected with sgRNAs as described above but without selection and GFP% was measured on day three (P0) and then every three days post-viral transduction until day 18 (P5). For CRISPR interference at the IL1 enhancer, SUIT2-empty or SUIT2-TP63 cells stably expressing catalytically-dead Cas9 fused with a KRAB repression domain in the Lenti-dCas9-KRAB-blast vector (addgene #89567) were infected in a pooled fashion with sgRNAs targeting IL1 enhancer regions (identified from TP63 and H3K27ac ChIP-seq in these lines) in the LRNG vector. Stable cell lines were selected by G418 selection for seven days prior to their use in conditioned media experiments. For doxycycline-inducible CRISPR based targeting of TP63 in T3M4 and KLM1 cells, Cas9 from the LentiV-Cas9-puro vector was cloned into the doxycycline-regulated TREtight-cDNA-EFS-rtTA-P2A-Puro vector (Somerville et al., 2018) to generate the YXP-Cas9-puro vector. T3M4 and KLM1 cells were infected with the YXP-Cas9-puro vector, selected with 3µg/ml puromycin to generate stable cell line before infection with sgRNAs targeting TP63 (sgTP63#4) or a control sgRNA (sgNEG) in LRNG vector. Two days post infection with sgRNAs, transduced cells were selected with 1mg/ml of G418 for three days and were subsequently treated with doxycycline (1µg/ml) for 48 hours prior to harvesting RNA and lysate for RNA-seq and western blot analysis, respectively. sgRNA sequences can be found in Table S5.

### RNA extraction and RT-qPCR analysis

Total RNA was extracted using TRIzol reagent following the manufacturer’s instructions. For RNA extraction from cells cultured in Matrigel, cells were lysed by adding TRIzol reagent directly to the Matrigel dome. 200ng-1µg of total RNA was reverse transcribed using qScript cDNA SuperMix (Quanta bio; 95048-500), followed by RT-qPCR analysis with Power SYBR Green Master Mix (Thermo Fisher Scientific; 4368577) on an ABI 7900HT fast real-time PCR system. Gene expression was normalized to *GAPDH* or *Gapdh* for human and murine gene expression analysis, respectively. RT-qPCR primers used can be found in Table S5.

### Cell Lysate Preparation and Western Blot Analysis

Cell cultures were collected and 1 million cells were counted by trypan blue exclusion and washed with ice cold PBS. Cells were then resuspended in 100µl PBS and lysed with 100µl of 2x Laemmli Sample Buffer supplemented with β-mercaptoethanol by boiling for 30 minutes. Samples were centrifuged at 4°C for 15 minutes at 10,000xg and the supernatant was used for western blot analysis with standard SDS-PAGE-based procedures. Primary antibodies used were TP63 (Cell Signaling; 39692), HSC70 (Santa Cruz; sc-7298), ACTB (Sigma-Aldrich; A3854) and Cas9 (Epigentek; A-9000-050) and proteins were detected using HRP-conjugated secondary antibodies.

### ELISA

For ELISA of plasma, blood was harvested via cardiac puncture from anaesthetized mice at end point and plasma separated by centrifugation. For ELISA of media, conditioned media was harvested as described above. ELISA assays used were IL-1α (DLA50; R&D Systems), IL-1β (DLB50; R&D Systems) and IL-6 (M6000B; R&D Systems).

### In Vitro Luciferase Imaging

To generate luciferase expressing mouse PSC cultures, cells were infected with a luciferase transgene in a Lenti-luciferase-blast vector (Somerville et al., 2018) and stable cell lines were generated by selection with 10µg/ml blasticidin. Cells were plated at a density of 10,000 cells per 20µl of Matrigel in each well of a black, clear-bottom, ultra-low attachment 96-well plate (10014-318; CELLSTAR) and 200µl of conditioned media was added. Cells were imaged following four days of culture using an IVIS Spectrum system (Caliper Life Sciences) six minutes post addition of D-Luciferin (150µg/ml) to each well. For luciferase-based proliferation assays of SUIT2-TP63 cells in *vitro*, parental SUIT2 cells were first infected with Lenti-luciferase-blast vector and stable SUIT2-luciferase cell lines were generated by selection with 10µg/ml blasticidin. SUIT2-luciferase cells were then infected with TP63 cDNA in LentiV-ΔNp63-neo vector (Somerville et al., 2018) or the empty vector as a control. Two days post infection, transduced cells were selected with 1mg/ml of G418 and on day seven post infection, cells were counted by trypan blue exclusion and seeded at a density of 500 cells per well in 200µl of media in black 96-well plates (137101; Thermo). Cells were imaged daily as described above.

### In Vivo Orthotopic Transplantations and Bioluminescence Imaging

All animal procedures and studies were approved by the Cold Spring Harbor Laboratory Animal Care and Use Committee in accordance to IACUC. Transplantation of human PDA cells and mouse pancreatic organoid cultures have been described previously (Boj et al., 2015; Somerville et al., 2018). For mM1 organoid transplantations, 1×10^5^ cells were prepared from organoid cultures as a 45µL suspension of 50% Matrigel in PBS and injected into the pancreas. For TP63 knockout experiments *in vivo*, KLM1-Cas9 cells were first infected with Lenti-luciferase-blast vector and a stable KLM1-Cas9-luciferase cell line was generated by selection with 10µg/ml blasticidin. These cells were subsequently infected with control or TP63 sgRNAs in LRNG vectors as described above. G418-selected cells were counted by trypan blue exclusion and 5×10^4^ viable cells were prepared as a 50µL suspension of 50% Matrigel in PBS and injected into the pancreas. For *in vivo* experiments using SUIT2-luciferase cells, these cells were infected with LentiV-ΔNp63-neo vector or the empty vector as a control and two days post infection, transduced cells were selected 1mg/ml of G418 for five days and on day seven post infection, G418-selected cells were counted by trypan blue exclusion and 2×10^4^ viable cells were prepared as a 50µL suspension of 50% Matrigel in PBS and injected into the pancreas. For bioluminescence imaging, mice were intraperitoneally (IP) injected with D-Luciferin (50mg/kg) and analyzed using an IVIS Spectrum system (Caliper Life Sciences) ten minutes post IP injection.

### Histology and Immunohistochemistry

Histological analysis and TP63 immunohistochemistry (IHC) was performed as previously described (Somerville et al., 2018). Primary antibodies used were TP63 (Cell Signaling; 39692). Hematoxylin and eosin and Masson’s trichrome staining were performed according to standard protocols. Stained sections were scanned with Aperio ScanScope CS and to quantify TP63, the percentage of strong positive nuclei was calculated relative to the total number of nuclei with the ImageScope nuclear v9 algorithm.

### Flow Cytometry and Cell Sorting

Tumors were processed as previously described (Öhlund et al., 2017). For sorting of cancer cells, fibroblasts and immune cells from xenografted tumors, cells were stained for 30 minutes with anti-mouse CD45-PerCP-Cy5-5 (103132; BioLegend), PDPN-APC/Cy7 (127418; BioLegend) and anti-human CD326 (EPCAM)-AlexaFluor 647 (324212; BioLegend), anti-human/mouse E-Cadherin-AlexaFluor 647 (147304; BioLegend) and DAPI for 15 minutes. Cells were sorted on the FACSAria cell sorter (BD) for DAPI/CD45^−^EPCAM/E-Cadherin^+^ (cancer cells), DAPI/CD45/EPCAM/E-Cadherin^−^PDPN^+^ (fibroblasts) and DAPI^-^CD45^+^ (immune cells) cell populations. Sorted cells were pelleted and resuspended in TRIzol for RNA extraction. For flow cytometric analysis of processed mM1 tumors, antibodies used for analysis of myeloid cells were: anti-mouse CD45-BV510 (103138; BioLegend), F4/80-BV785 (107645; BioLegend), Ly6G/Ly6C (Gr-1)-PE (108408; BioLegend) and CD11b-PE-Cy7 (101216; BioLegend); antibodies used for analysis of T-cells cells were: anti-mouse CD45-BV510 (103138; BioLegend), CD8a-APC-Cy7 (100714; BioLegend), CD4-APC (100516; BioLegend) and TCRβ-PE-Cy7 (109222; BioLegend). For flow cytometric analysis of processed xenografted tumors, antibodies used were anti-mouse CD45-PerCP-Cy5.5 (103132; BioLegend), F4/80-BV785 (107645; BioLegend), Ly6G-Ly6C (Gr-1)-PE (108408; BioLegend) and CD11b-AlexaFluor 488 (101217; BioLegend). Fixable viability dye eFluor450 (eBioscience) was used to differentiate between live and dead cells.

### Neutrophil Isolation and Transwell Migration Assays

Neutrophil isolation was performed as previously described (Albrengues et al., 2018). Transwell migration assays were performed with conditioned media derived from SUIT2-empty or SUIT2-TP63 cells cultured alone or in combination with mouse PSCs for five days in Matrigel. 5×10^3^ SUIT2 cells and 4×10^4^ PSCs were used for the co-cultures. The conditioned media was added to the lower chamber, and primary mouse neutrophils (2.5×10^5^) were seeded in 0.5% FBS containing DMEM in the upper chamber of a 3µm FluoroBlok cell culture insert (08-772-141; Corning) in a 24 well plate. 500ng of recombinant mouse (250-11; Peprotech) or human (300-11; Peprotech) CXCL1 was added to DMEM with 5% FBS on the day of seeding neutrophils as controls. After 24 hours, the FluoroBlok membrane was stained with DAPI (0.05mg/ml; D1306; Thermo Fisher Scientific) for 5 minutes, rinsed in water and mounted onto glass slides using mounting media (17985-16; Electron Microscopy Sciences). The number of invading neutrophils was counted in 10 random fields of view using a fluorescence microscope (Leica SP8 Confocal microscope).

### RNA-seq Library Construction

RNA-seq libraries were constructed using the TruSeq sample Prep Kit V2 (Illumina) according to the manufacturer’s instructions. Briefly, 0.5-2µg of purified RNA was poly-A selected and fragmented with fragmentation enzyme. cDNA was synthesized with Super Script II Reverse Transcriptase (Thermo Fisher; 18064014), followed by end repair, A-tailing and PCR amplification. RNA-seq libraries were single-end sequenced for 50bp, or for xenograft RNA-seq analysis paired-end sequenced for 150bp, using an Illumina NextSeq platform (Cold Spring Harbor Genome Center, Woodbury).

### RNA-Seq Data Analysis

Single end 50bp sequencing reads were mapped to the mm10 or hg38 genomes using HISAT2 with standard parameters (Kim et al., 2015). Structural RNA was masked and, unless stated otherwise, differentially expressed genes were identified using Cuffdiff (Trapnell et al., 2010). All the following analysis was performed on genes with an RPKM value no less than 2 in either control or experimental samples. Heat maps of standardized expression values for differentially expressed genes were generated using Morpheus from the Broad Institute (https://software.broadinstitute.org/morpheus). To generate ranked gene lists for Pre-ranked GSEA, genes were ranked by their mean log_2_ fold change between the two experimental groups of interest. Gene ontology analysis was performed using Metascape (Zhou et al., 2019). For clustering of mouse PSC cultures, the log_2_(RPKM+1) values for each cell line was used to generate a heat map of similarity matrix by Pearson correlation, which was subsequently clustered by Euclidean distance with average linkage using Morpheus. Group 1 and Group 2 cultures were then analysed using Cuffdiff and those genes with log_2_ fold change >1 and a FDR value <0.05 were considered and differentially expressed. For GSEA using the Broad Institutes Hallmark gene sets, mouse gene symbols were converted to their orthologous human symbols and ranked by their average log_2_ fold change in the respective cultures. For RNA-seq analysis of flow sorted cancer cells, fibroblasts and immune cells, those genes with a FDR value <0.05 when comparing sgTP63#4 versus sgNEG tumors were considered and differentially expressed. For xenograft RNA-seq analysis, paired end 150bp sequencing reads were mapped to the mm10 or hg38 genomes using HISAT2 with standard parameters. The alignment sam files were converted to bam format using the samtools view command. To disambiguate between the human and mouse reads, Disambiguate (Ahdesmäki et al., 2016) was run on the bam files, and the unambiguous reads were used for RNA quantification and differential expression analysis using DESeq2 (Love et al., 2014). For human transcripts, those genes with log_2_ fold change >1 and a FDR value <0.1 were considered as differentially expressed. For mouse transcripts, those genes with a FDR value <0.1 were considered as differentially expressed. To generate a ranked list of mouse genes for GSEA analysis genes the bottom 30% of expressed genes across all groups were removed. The iCAF and myCAF gene signatures were defined as the top 200 up- and down-regulated genes in iCAFs versus myCAFs respectively from the study by Öhlund et al (2017). The Squamous-PDA Identity signature has been previously described (Somerville et al., 2018). RNA-seq datasets following TP63 expression in SUIT2, PATU8988S, HPAFII, AsPC1 and CFPAC1 cells as well as TP63 knockout or knockdown in BxPC3 and hF3 cells were from a previous study (Somerville et al., 2018).

### ChIP-Seq Library Construction and Analysis

ChIP procedures were performed as previously described (Somerville et al., 2018). The antibody used for ChIP-seq in this study was H3K27ac (ab4729; abcam). ChIP-seq libraries were constructed using Illumina TruSeq ChIP Sample Prep kit following manufacture’s protocol. Briefly, ChIP DNA was end repaired, followed by A-tailing and size selection (300-500bp) by gel electrophoresis using a 2% gel. 15 PCR cycles were used for final library amplification which was analyzed on a Bioanalyzer using a high sensitivity DNA chip (Agilent). ChIP-seq libraries were single-end sequenced for 50bp using an Illumina NextSeq platform (Cold Spring Harbor Genome Center, Woodbury). Single end 50bp sequencing reads were mapped to the hg19 genome using Bowtie2 with default settings (Langmead and Salzberg, 2012). After removing duplicated mapped reads using SAM tools (Li et al., 2009), MACS 1.4.2 was used to call peaks using input genomic DNA as control (Feng et al., 2012). Only peaks enriched greater than or equal to 10-fold over input samples were used for subsequent analyses. Metagene plots were made by centering H3K27ac ChIP-seq regions on previously defined Squamous Elements (Somerville et al., 2018) extended to ±10,000bp with 100bp bins. Super enhancer analysis was performed using Rank Ordering of Super-Enhancers (ROSE) as described (Lovén et al., 2013; Whyte et al., 2013). TP63 ChIP-seq in BxPC3 cells and H3K27ac ChIP-seq in hN34, hN35, hT85, PATU8988S, AsPC1, HPAFII, SUIT2, hF3 and BxPC3, as well as H3K27ac in SUIT2 or BxPC3 cells following TP63 expression or knockout were from a previous study (Somerville et al., 2018).

### Statistics

For graphical representation of data and statistical analysis, GraphPad Prism was used. Statistical analysis was performed as described in the figure legends.

## Supplemental material

Table S1 contains the genes corresponding to the Group 1 and Group 2 PSC clusters and the ranked gene list used for GSEA. Tables S2 contains the iCAF and myCAF gene signatures. Table S3 contains genes significantly upregulated in the human and mouse compartments of SUIT2-TP63 versus SUIT2-empty tumors and ranked gene lists used for GSEA. Table S4 contains genes significantly downregulated in each sorted fraction of TP63-negative versus TP63-positive KLM1 tumors and gene lists used for GSEA. Table S5 contains RT-qPCR primer sequences and sgRNA sequences used in this study.

## Acknowledgements

The authors would like to thank the Cold Spring Harbor Cancer Center Support Grant (CCSG) shared resources: Bioinformatics Shared Resource, Next Generation Sequencing Core Facility, R. Rubino and J. Habel Animal Resources for technical assistance with the mouse transplantation assays, P. Moody and C. Viola in the Flow Cytometry Facility, Animal and Tissue Imaging, and the Animal Facility. T.D.D.S. was supported by a grant from the State of New York, contract no. C150158. C.R.V. was supported by Pershing Square Sohn Cancer Research Alliance, the Cold Spring Harbor Laboratory and Northwell Health Affiliation, the National Cancer Institute (NCI) 5P01CA013106-Project 4 and 1RO1CA229699, and a Career Development Award from the Pancreatic Cancer Action Network-American Association for Cancer Research (AACR) 16-20-25-VAKO. This work was supported by the Lustgarten Foundation, where D.A.T is a distinguished scholar and Director of the Lustgarten Foundation–designated Laboratory of Pancreatic Cancer Research. D.A.T is also supported by the Cold Spring Harbor Laboratory Association and the National Institutes of Health (NIH 5P30CA45508, 5P50CA101955, P20CA192996, U10CA180944, U01CA210240, U01CA224013, 1R01CA188134, and 1R01CA190092). C.R.V., D.A.T., and M.E. were supported by the Thompson Family Foundation and Simons Foundation. In addition, we are grateful for support from the following: the Human Frontiers Science Program (LT000195/2015-L for G.B.), EMBO (ALTF 1203-2014 for G.B.), A Deutsche Forschungsgemeinschaft (DFG) Research Fellowship (DA 2249/1-1 for J.D.P).

---

**Figure S1.**
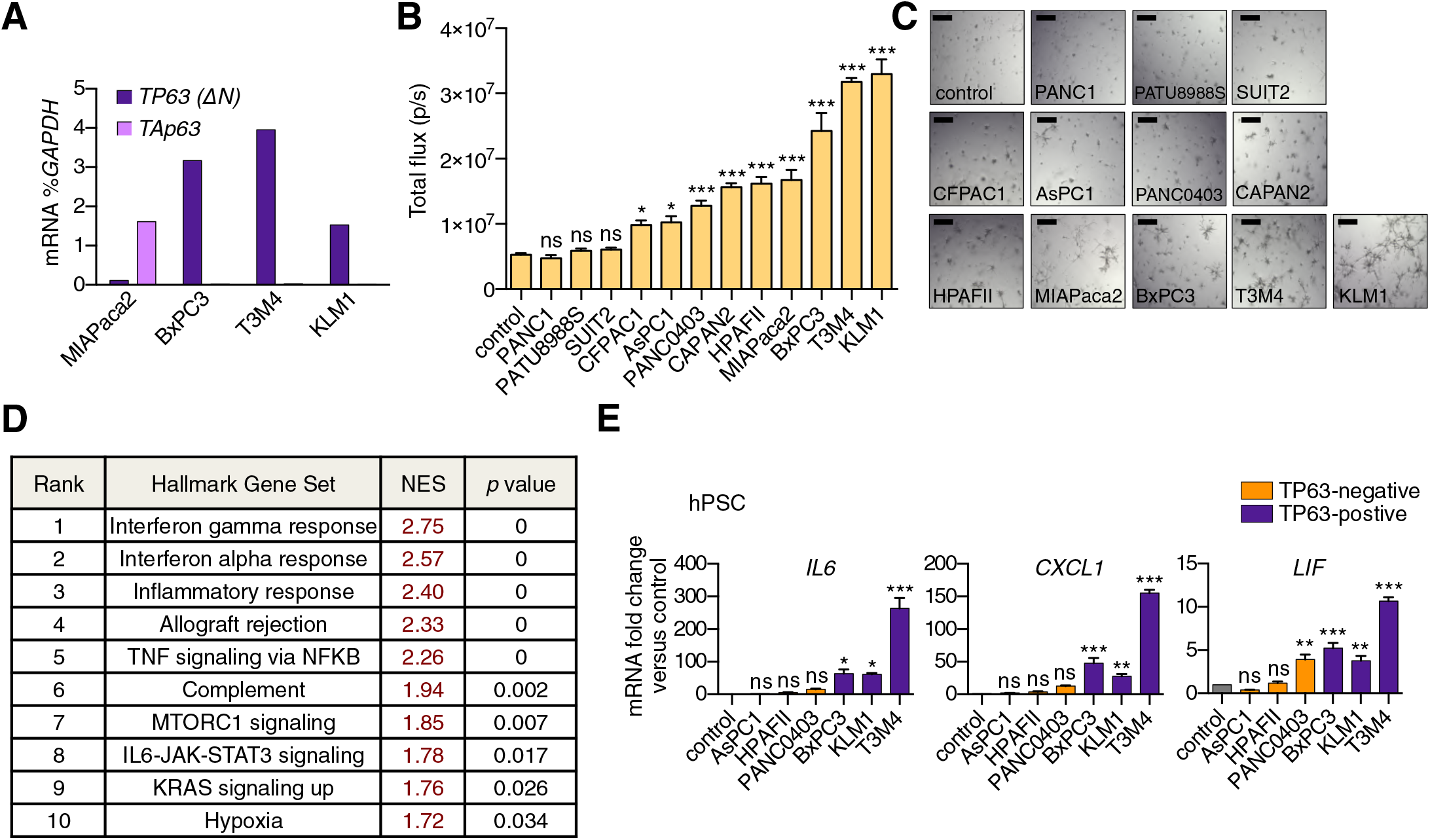
A secretory phenotype of TP63-positive PDA cells that promotes inflammatory gene expression changes in CAFs *in vitro*. Related to Figure 1. (A) Bar chart shows RT-qPCR analysis for *TP63* isoforms in MIAPaca2, BxPC3, T3M4 and KLM1 PDA cell lines. (B) Luciferase-based quantification of mouse PSC proliferation in Matrigel following four days of culture with conditioned media from the indicated human PDA cell lines. Mean+SEM is shown. n=5 technical repeats (C) Representative bright-field images from (B). Scale bar indicates 200µm. (D) Table summarizing GSEA evaluating Hallmark gene sets for their enrichment in Group 2 versus Group 1 cultures. Gene sets with a FWER *p* value <0.05 are shown. (E) Bar chart showing RT-qPCR analysis for iCAF markers (*IL6*, *CXCL1*, *LIF)* following culture of human PSCs in Matrigel for four days with conditioned media from the indicated human cell lines. Mean+SEM is shown. n=3. ***p <0.001, **p <0.01, *p <0.05 by one-way ANOVA with Dunnett’s test for multiple comparisons. ns, not significant.

**Figure S2.**
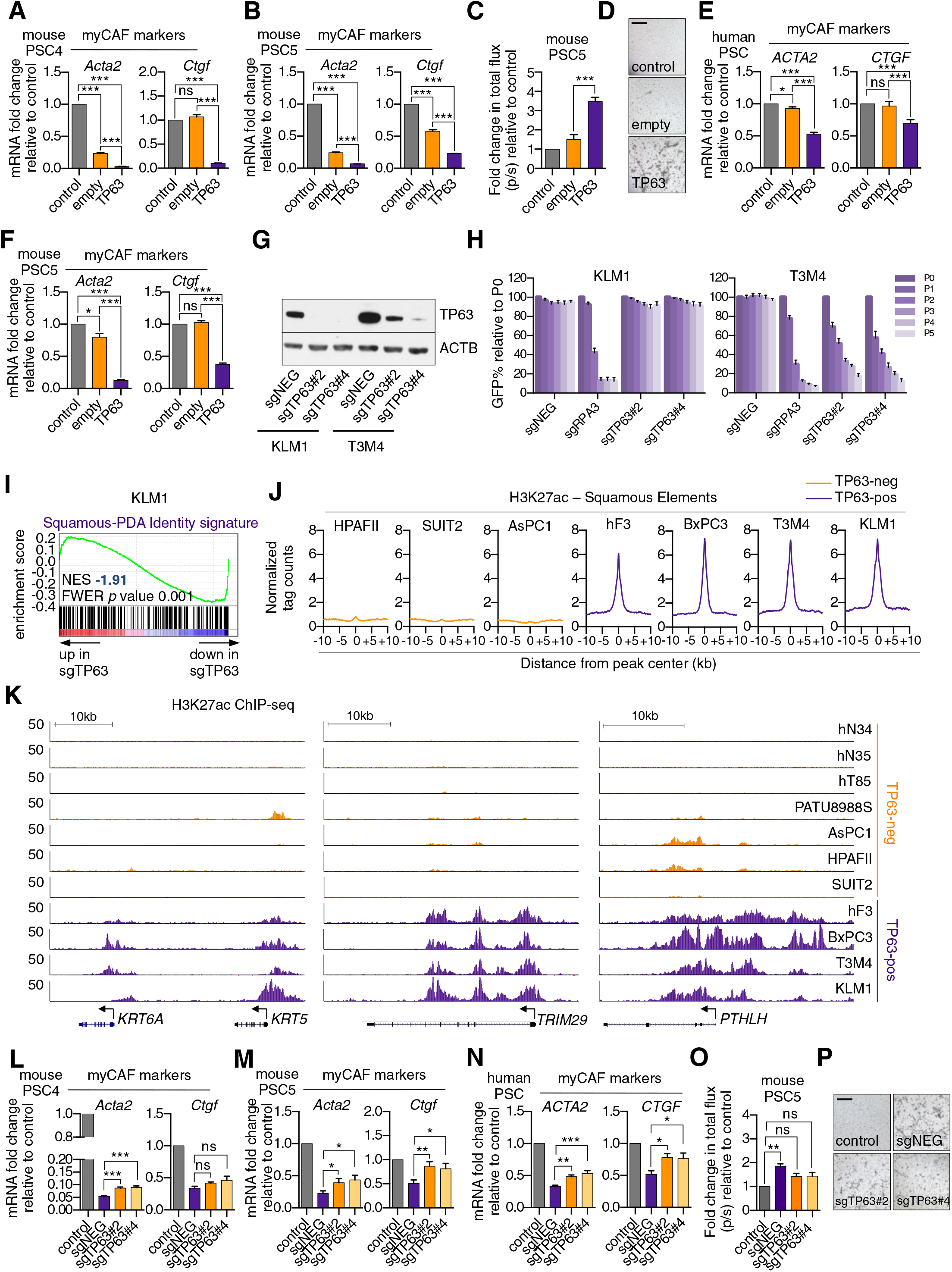
TP63 expression in PDA cells drives a secretory phenotype that induces iCAF formation in vitro. Related to Figure 2. (A-B) Bar charts showing RT-qPCR analysis for myCAF markers (*Acta2*, *Ctgf*) following culture of the two mouse PSC lines in Matrigel for four days with conditioned media from SUIT2-empty or SUIT2-TP63 cells. (C) Luciferase-based quantification of mouse PSC proliferation in Matrigel. (D) Representative bright-field images from (C). Scale bar indicates 500µm. (E) Bar charts showing RT-qPCR analysis for myCAF markers (*ACTA2*, *CTGF*) following culture of human PSCs in Matrigel for four days with conditioned media from SUIT2-empty or SUIT2-TP63 cells. (F) Bar chart shows RT-qPCR analysis for myCAF markers (*Acta2*, *Ctgf*) following culture of mouse PSCs in conditioned media from mM1-empty or mM1-TP63 cells. (G) Representative western blot analysis for ACTB and TP63 following TP63 knock out in KLM1-Cas9 and T3M4-Cas9 cells. (H) Competition based GFP depletion assays of KLM1-Cas9 cells (left panel) or T3M4-Cas9 cells (right panel) following infection with the indicated sgRNAs linked to GFP. (I) GSEA plot evaluating the Squamous-PDA Identity signature following doxycycline-inducible knock out of TP63 and subsequent RNA-seq analysis KLM1 cells. The Squamous-PDA Identity signature was defined previously (Somerville et al., 2018). (J-K) Metagene representation (J) and browser track examples (K) of H3K27ac signal at Squamous Elements in TP63-negaive and TP63-postive PDA cells. Normal organoids: hN34, hN35; PDA organoids: hF3, hT85; PATU: PATU8988S. (L-N) Bar charts showing RT-qPCR analysis for myCAF marker genes following culture of two mouse PSC cell lines (L and M) or human PSCs (N) in Matrigel for four days with conditioned media from KLM1-Cas9 cells infected with the indicated sgRNA. (O) Luciferase-based quantification of mouse PSC proliferation in Matrigel cultured with conditioned media from KLM1-Cas9 cells infected with sgRNAs targeting TP63 or control (sgNEG). (P) Representative bright-field images from (O). Scale bar indicates 500µm. For all experiments control media represents DMEM supplemented with 5% FBS. Matrigel only represents conditioned media from Matrigel cultured with control media only. Mean+SEM is shown. n=3. For A, B, E and F, *** p < 0.001, ** p < 0.01, * p < 0.05 by Student’s t-test and for C, *** p < 0.001, ** p < 0.01, * p < 0.05 by one-way ANOVA with Dunnett’s test for multiple comparisons. ns, not significant.

**Figure S3.**
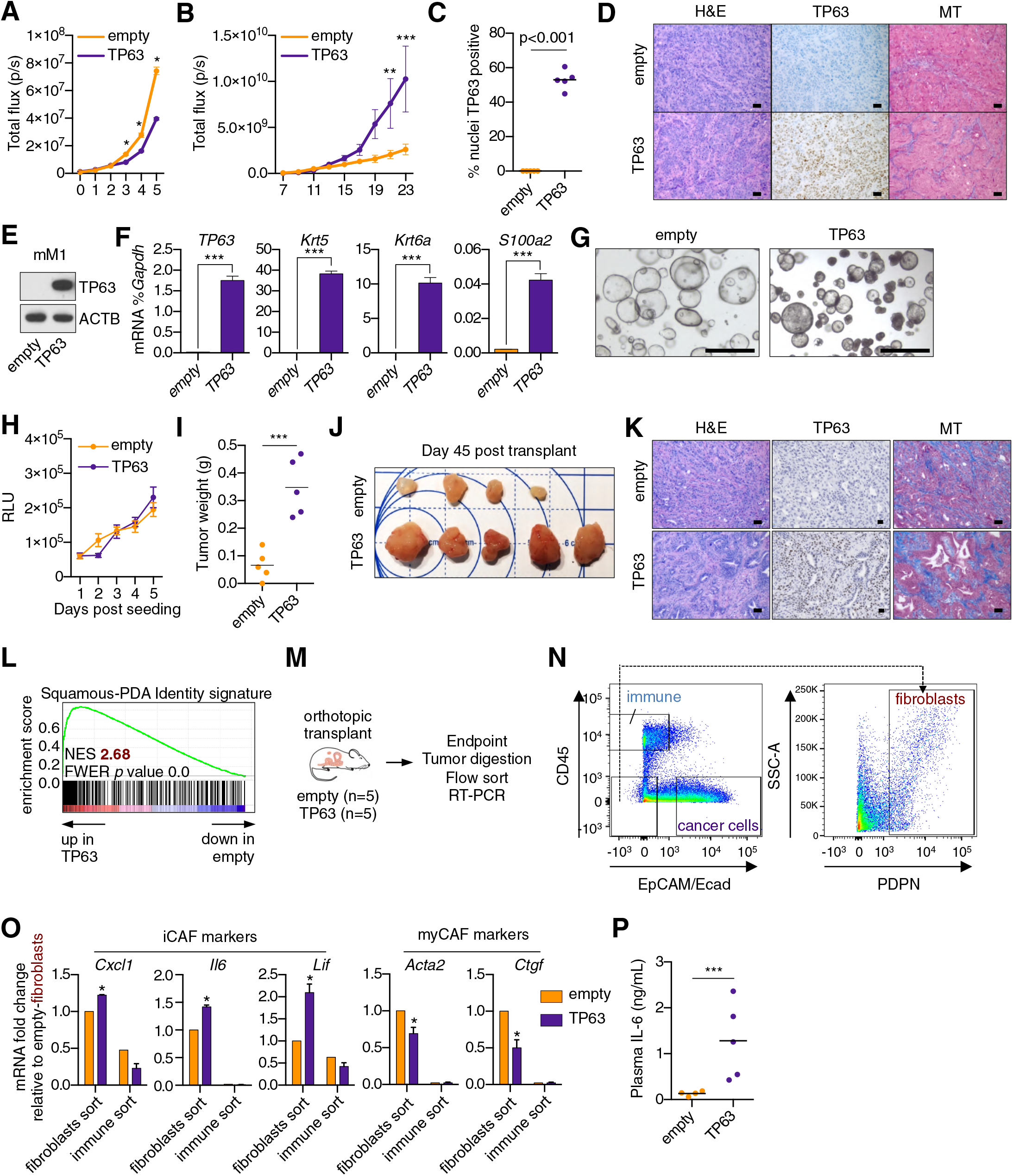
Ectopic expression of TP63 in PDA cells promotes inflammation-associated transcriptional changes in the tumor microenvironment. Related to Figure 3. (A) Luciferase-based quantification of SUIT2-empty and SUIT2-TP63 cell proliferation *in vitro*. Mean±SEM values are shown. n=6 technical repeats. (B) Quantification of the bioluminescence signal following transplantation of SUIT2-empty or SUIT2-TP63 cells to the pancreas of NSG mice. Mean±SEM in shown. Mice were imaged every two days between day seven and 23 post-transplantation. n=5 mice per group. (C) Quantification of TP63 expression by immunohistochemical staining of tumors. *p* value determined by Student’s t-test. (D) Representative images from (C). MT, Masson’s trichrome. Scale bar indicates 50µm. (E) Representative western blot analysis for ACTB and TP63 in mM1-empty or mM1-TP63 cells. (F) Bar chart showing RT-qPCR analysis for squamous markers (*TP63*, *Krt5*, *Krt6a* and *S100a2*) in mM1-empty and mM1-TP63 cells. Mean+SEM is shown. n=3. (G) Representative bright-field images of mM1 organoid cultures. (H) Quantification of *in vitro* cell growth measured by CellTiter-Glo. RLU, relative luminescence units. (I-J) Quantification of tumor volume (I) and images of tumors (J) on day 45 post-transplantation of mM1 orgaoinds to the pancreas of C57BL/6 mice. ***p<0.001 by Student’s t-test. (K) Representative images from (J). MT, Masson’s trichrome. Scale bar indicates 50µm. (L) GSEA plot evaluating the Squamous-PDA Identity signature (Somerville et al., 2018) within the human cancer cell compartment of TP63-positive versus negative tumors. (M-O) RT-qPCR analysis of flow-sorted tumor samples. (M) Schematic of experimental workflow. (N) Representative flow cytometry plots showing the gating strategy for enriching human cancer cells, mouse fibroblasts and mouse immune cells. (O) Bar chart showing RT-qPCR analysis for the indicated genes in the indicated sorted fraction and tumor sample. \*\*\**p* < 0.001, \*\**p* < 0.01, \**p* < 0.05 by two-way ANOVA with Sidak’s test for multiple comparisons. (P) ELISA of IL-6 in the plasma of mice at end point. n=4-5 mice per group. ***p < 0.001 by Student’s t-test.

**Figure S4.**
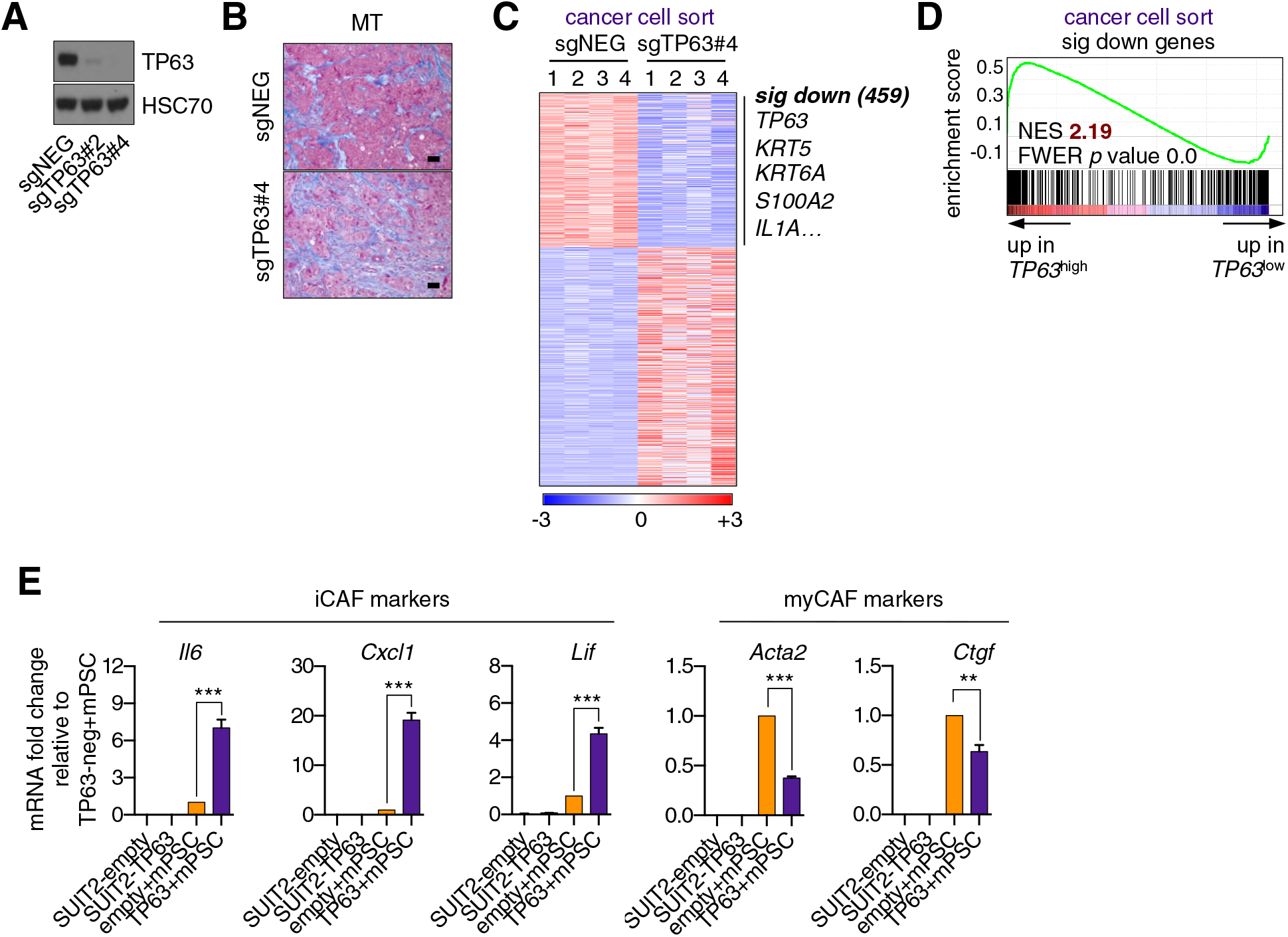
Knockout of TP63 in an orthotopic PDA tumor model attenuates the inflammatory signature of fibroblasts and immune cells. Related to Figure 4. (A) Representative western blot analysis for HSC70 and TP63 in the KLM1-Cas9 cells following TP63 knock out with CRISPR. (B) Representative images from Fig. 4D. Scale bar indicates 50µm. MT, Masson’s trichrome. (C) Heatmap representation of differentially expressed genes in the human cancer cell sorted fraction. (D) GSEA plot evaluating the 459 significantly down-regulated human genes based on their expression in *TP63*^high^ versus *TP63*^low^ PDA patient samples from the study by Bailey et al (2016). Patients were designated as *TP63*^high^or *TP63*^low^ as described in Somerville et al (2018). (E) Bar charts show RT-qPCR analysis for the iCAF markers (*Il6*, *Cxcl1*, *Lif*; left panel) and myCAF markers (*Acta2*, *Ctgf*; right panel) in the indicated culture conditions. Mean+SEM is shown. n=3. ***p < 0.001, **p < 0.01 by Student’s t-test.

**Figure S5.**
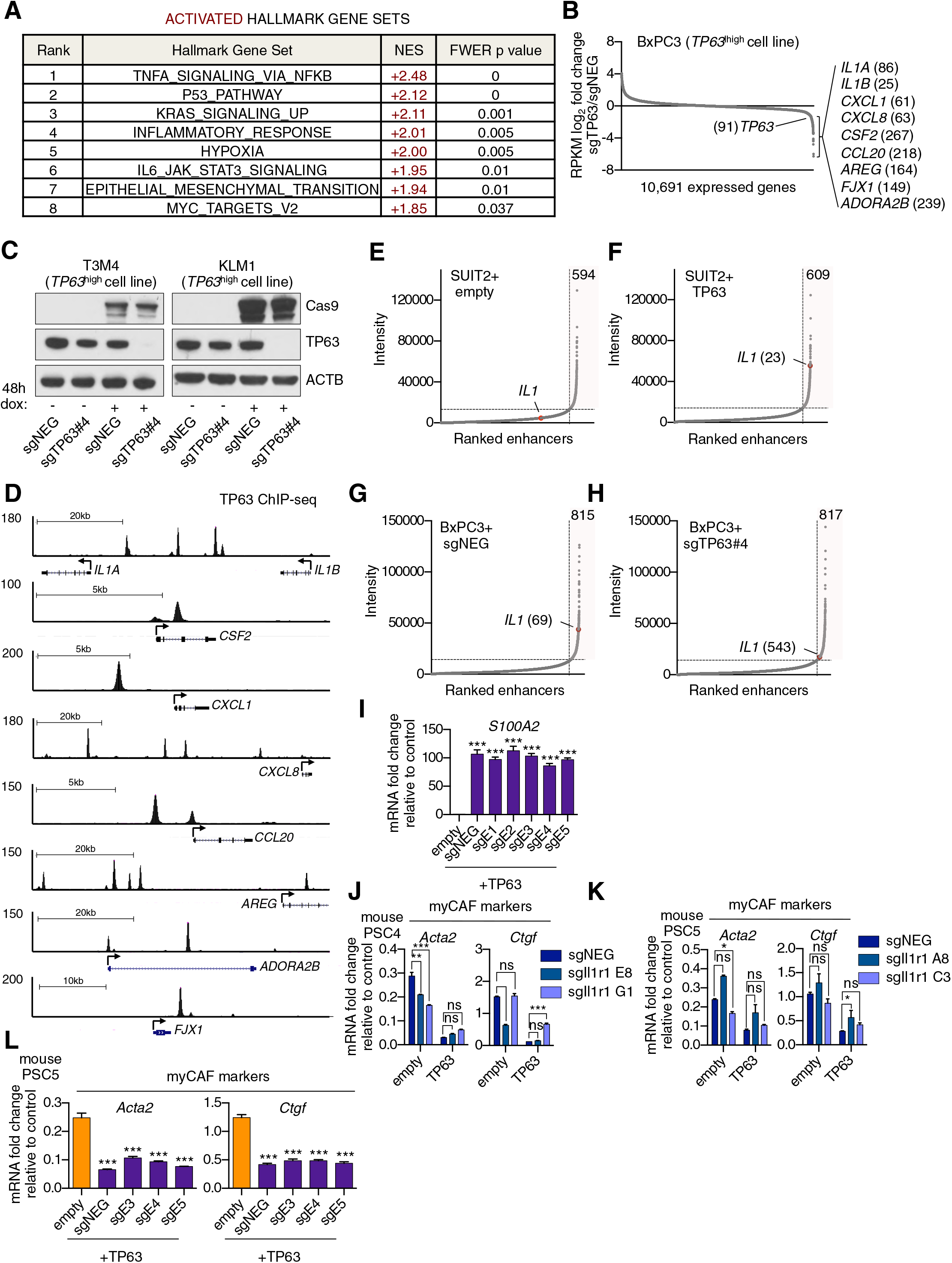
TP63 activates enhancer elements and transcription of genes encoding pro-inflammatory cytokines in PDA cells. Related to Figure 5. (A) Table shows evaluation of the indicated Hallmark gene sets by GSEA upon TP63 expression in SUIT2 cells. Gene sets are ranked by their NES score and only those with a FWER *p* value <0.05 are shown. (B) Scatter plot shows the mean log_2_ fold change of expressed genes upon TP63 knock out in BxPC3 cells. Genes with a mean log_2_ fold change >1 in this dataset and the SUIT2-TP63 dataset (Fig. 5C) that are also found in the gene signatures shown in Fig. 5A and Fig. 5B are highlighted along with the rank. Data are from Somerville et al (2018). (C) Representative western blot analysis for ACTB, TP63 and Cas9 in KLM1 and T3M4 cells lentivirally transduced with a doxycycline-inducible Cas9 vector and the indicated sgRNA. (D) ChIP-seq profiles of TP63 in BxPC3 cells surrounding the genes shown in Fig. 5D. (E-H) Scatter plots showing enhancers ranked by intensity in SUIT2-empty (E), SUIT2-TP63 (F) and BxPC3-Cas9 cells transduced with a control sgRNA (G) or an sgRNA targeting TP63 (H). Shaded box highlights the number of super enhancers identified and the super enhancer associated with the IL1 locus is highlighted along with its rank. (I) RT-qPCR analysis for *S100A2* in SUIT2-empty or SUIT2-TP63 cells infected with dCas9 fused with the KRAB repression domain and the indicated sgRNAs. Mean+SEM is shown. n=3. *** p < 0.001, ** p < 0.01, * p < 0.05 versus empty control by one-way ANOVA with Dunnett’s test for multiple comparisons. (J-K) Bar charts showing RT-qPCR analysis for the indicated myCAF markers (*Acta2*, *Ctgf*) following culture of two mouse PSC lines from which the IL1 receptor was clonally knocked out with CRISPR in Matrigel for four days with conditioned media from SUIT2-empty or SUIT2-TP63 cells. (L) Bar charts showing RT-qPCR analysis for myCAF markers (*Acta2*, *Ctgf*) following culture of PSCs in Matrigel for four days with the conditioned media harvested from cells shown in Fig. 5I. Mean+SEM is shown. n=3. ***p <0.001, **p <0.01, *p <0.05 by one-way ANOVA with Dunnett’s test for multiple comparisons.

